# IL-18 metabolically reprograms CAR-expressing natural killer T cells and enhances their antitumor activity against neuroblastoma

**DOI:** 10.1101/2025.04.18.649551

**Authors:** Gabriel A. Barragán, David A. de la Cerda, Elisa Landoni, Kshiti Dholakia, Piotr Humeniuk, Leidy D. Caraballo, Ying Wang, Gengwen Tian, Boning Yang, Linjie Guo, Michael S. Wood, Xavier Rios, Xin Xu, Amy N. Courtney, Erica J. Di Pierro, Gianpietro Dotti, Leonid S. Metelitsa

**Affiliations:** Baylor College of Medicine; University of North Carolina, Chapel Hill

**Keywords:** IL-18, IL-15, CAR, Natural Killer T cell, Metabolism, Neuroblastoma

## Abstract

Invariant natural killer T cells (NKTs) have intrinsic anti-tumor properties that make them promising candidates for chimeric antigen receptor (CAR)-based immunotherapies. Transgenic cytokine expression has been shown to enhance the potency of cellular immunotherapies, and we hypothesized that co-expressing IL-18 alone or with IL-15 would boost CAR-NKT therapeutic potential. To test this hypothesis, we generated retroviral constructs expressing IL-15 and/or IL-18 with the inducible caspase 9 (iC9) safety switch and co-transduced them with a GD2-specific CAR into human NKTs. Co-expression of IL-18 or IL-15/IL-18 increased GD2.CAR-NKT *in vitro* cytotoxicity, proliferation, and cytokine secretion compared to IL-15 alone. In a metastatic neuroblastoma model, GD2.CAR-NKTs expressing constructs with IL-18 controlled tumor growth better than cells expressing IL-15 only, but mice in the IL-15/IL-18 group developed severe toxicities not observed in the IL-18-only group. Mechanistically, we found that IL-18 drives a distinct transcriptional profile from IL-15 in CAR-NKTs marked by lower expression of exhaustion gene signatures and enrichment of metabolism-related processes. Finally, targeted metabolomics revealed that IL-18 induces broad metabolic reprogramming in CAR-NKTs including enhancement of oxidative phosphorylation, glycolysis, glutaminolysis and purine metabolism. These results support the use of IL-18 in developing the next generation of cytokine-armed CAR-NKT cancer immunotherapy.

## Introduction

Despite significant progress in treating hematologic malignancies, T cells expressing chimeric antigen receptors (CARs) have generated few clinical responses in patients with solid tumors to date.^1^ This is partially because CAR-T cells do not localize well to tumor sites and do not have the metabolic fitness to avoid rapid exhaustion and loss of function in the immunosuppressive solid tumor microenvironment (TME).^2^ Strategies to enhance the trafficking of CAR-T cells to tumors and boost therapeutic effector cell function in the TME are urgently needed to achieve durable control of solid tumors while managing toxicity.

Compared to conventional T cells, several types of innate or innate-like lymphocytes including NK, γ/δ T, and CD1d-restricted invariant natural killer T cells (NKTs) have intrinsic antitumor properties including the ability to effectively home to tumor sites. ^3–6^ These unconventional effectors can also be redirected using tumor antigen-specific CARs to effectively target solid malignancies.^7–10^ We recently found that despite showing similar levels of cytotoxicity to CAR-T cells *in vitro*, CAR-NKTs exert significantly more potent *in vivo* antitumor activity in multiple solid tumor models.^11^ NKTs can also mediate indirect antitumor activity by killing or inhibiting M2-like tumor associated macrophages (TAMs) in a CD1d-dependent manner.^12,13^ In patients, NKT infiltration into primary tumors has been associated with improved outcomes in children^3^ and adults with cancer.^14^ These properties make NKTs a promising cellular platform for CAR-redirected cancer immunotherapy.

Neuroblastoma (NB) is a common extra-cranial solid tumor of childhood, and patients with high risk/relapsed disease have a five-year survival rate of between 20% and 50%.^15,16^ The GD2 ganglioside is expressed at high levels in NB cells and minimally expressed in normal tissues, making it a suitable target for GD2-specific immunotherapy with two FDA-approved anti-GD2 monoclonal antibodies now available.^17,18^ In the cellular immunotherapy space, several early-stage clinical trials testing GD2-specific CAR-T cells^19,20^ and NKT cells^21^ have demonstrated safety and evidence of antitumor activity. On the GINAKIT2 trial, NKTs were engineered to co-express a GD2.CAR and IL-15, a cytokine that has been shown to support NKT and CAR-NKT persistence and antitumor activity in preclinical tumor models.^22^ Beyond IL-15, the membrane-bound form of proinflammatory cytokine IL-12 has been shown to potently enhance GD2.CAR-NKT persistence and antitumor activity in xenogeneic tumor models.^23^ Like IL-12, IL-18 is a proinflammatory cytokine that induces effector programs in T and NK cells^24^ and stimulates IFNγ production by and antitumor activity of murine NKTs.^25,26^ IL-18 has already been shown to enhance CAR-T antitumor potential,^27–32^ providing rationale for ongoing phase 1 trials evaluating IL-18-armed CD19.CAR-T and GD2.CAR-T cells in patients with relapsed/refractory lymphomas (NCT04684563) and GD2-positive solid cancers (WWU19_0008), respectively.

Here, we studied the impact of IL-18 expression on human NKTs and CAR-NKTs to determine the therapeutic utility of this cytokine in this unique effector subset. We report that freshly isolated primary human NKTs exhibited a higher level of IL18Rα expression than NK and T cells from the same donors, and that IL-18 more potently enhanced CAR-NKT *in vivo* antitumor activity than IL-15 without increasing toxicity. Mechanistically, we demonstrated that IL-18 reprograms NKT cell metabolism, leading to improved functional fitness and resistance to exhaustion. In all, our results suggest that IL-18 expression is a promising strategy to improve the therapeutic potential of CAR-NKTs for neuroblastoma and potentially other solid tumors.

## Results

### Generation of CAR-NKT cells co-expressing IL-18 and/or IL-15 with an iCaspase 9 safety switch

While IL18Rα expression has been demonstrated in cord blood-expanded CD4^+^ NKT cells and adult healthy donor NKTs,^33,34^ a comparison of IL18Rα expression levels in NKTs versus other lymphocyte subsets has not been performed. To address this, we measured IL18Rα expression in freshly isolated peripheral blood mononuclear cells (PBMCs) by flow cytometry. Interestingly, NKT cells showed significantly higher IL18Rα levels than both T and NK cells (**Fig. 1A**), suggesting that NKT cells are responsive to IL-18. Additionally, IL-18Rα expression was maintained at a high level in NKTs following both primary and secondary stimulations with αGalCer-loaded antigen presenting cells (**Fig. 1B**).

**Figure 1.**
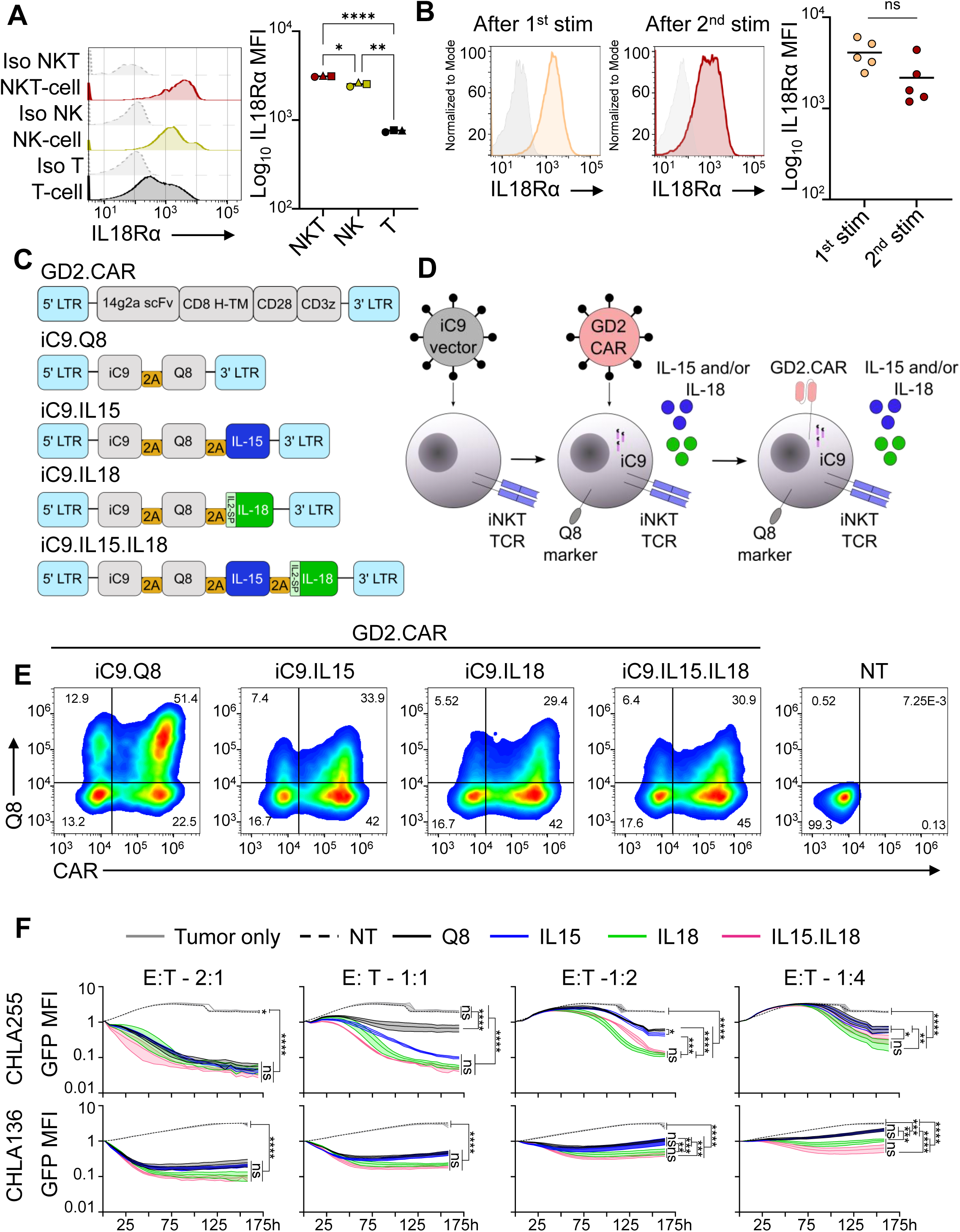
Generation of CAR-NKT cells co-expressing IL-15, IL-18, or IL-15/IL-18 with the iC9 safety switch. **A.** Flow cytometry histograms depicting IL18Rα expression in PBMC cell subsets from a healthy donor (left). Summary data of IL18Rα expression in PBMCs from *n*=3 healthy donors (right). * *p* < 0.05, ** *p* < 0.01, **** *p* < 0.0001, paired one-way ANOVA**. B.** Flow cytometry histograms depicting IL18Rα expression measured in NKT cells seven days after primary stimulation and seven days after secondary stimulation in the same donor. Grey histograms represent isotype control (left). Summary data of IL18Rα expression from *n*=5 donors. Mean is shown, ns=not significant, paired two-tailed t-test (right). **C.** Retroviral plasmids encoding GD2.CAR and iC9/cytokine constructs used in this study. LTR=long terminal repeat, scFv=single chain variable fragment, H=hinge, TM=transmembrane domain, IL2SP=IL2 signal peptide. **D.** CAR and iC9/cytokine construct co-transduction strategy, six days after isolation and primary stimulation, NKT cells were transduced with iC9/cytokine encoding retroviruses, 24 hours later cells were harvested and transduced with GD2.CAR retroviral supernatant. **E.** Two days after GD2.CAR transduction, co-transduced NKTs were stimulated using αGalCer loaded aAPCs and expanded for 10 days. Then, transduction rate was assessed by flow cytometry analysis using antibodies specific to the GD2.CAR and Q8 marker. Representative flow cytometry analysis of CAR/iC9 construct co-transduction efficiency 10 days after GD2.CAR transduction. **F.** GFP expressing CHLA255 (GD2^high^, top) and CHLA136 (GD2^int^, bottom) NB lines were co-cultured with CAR-NKTs co-transduced with the indicated constructs at different effector-to-target ratios. To determine tumor cytotoxicity, changes in GFP fluorescence were measured over the indicated period using the Incucyte system. Shown are results from a representative NKT donor of four with technical triplicates per condition. *ns*= not significant, * *p* < 0.05, ** *p* < 0.01, *** *p* < 0.001, **** *p* < 0.0001, one-way ANOVA.

Next, to explore the impact of IL-18 expression in CAR-NKTs versus clinically tested IL-15, we generated retroviral constructs encoding IL-18 and/or IL-15, the inducible Caspase 9 (iC9) safety switch,^35^ and the Q8 transduction marker (**Fig. 1C**).^36^ In these constructs, the IL-18 pro-peptide was replaced with the IL-2 signal peptide sequence to ensure production of biologically active and secreted IL-18.^16^ NKT cells were isolated from healthy donor PBMCs and expanded as previously described^37^ followed by transduction with iC9.IL15, iC9.IL18, iC9.IL15.IL18, or iC9.Q8 control and the GD2.CAR construct 24 hours later (**Fig. 1D**). The cells were then restimulated and cultured in the presence of IL-2 for 9-12 days, and transduction rate and NKT purity were assessed by flow cytometry (**Fig. 1E, Fig. S1A**). Co-expression of the iC9.IL18 or iC9.IL15.IL18 constructs affected NKT proliferation minimally during manufacturing when compared to cells expressing iC9.Q8 or iC9.IL15 (**Fig. S1B**), and NKTs expressing these constructs secreted their respective transgenic cytokines after stimulation with NB tumor cells (**Fig. S1C**). We confirmed activation of IL-15 signaling using intracellular staining for phosphorylated STAT5 and IL-18 signaling using an NFκB reporter construct (**Fig. S1D, E**).^38,39^

Addition of the chemical inducer of dimerization (CID) rimiducid (AP1903) effectively eliminated Q8^+^ cells, confirming the functionality of the iC9 safety switch (**Fig. S1F, G**). Next, we characterized CAR-mediated *in vitro* cytotoxicity by coculturing CAR-NKTs with CHLA255 (GD2^high^) or CHLA136 (GD2^int^) NB cell lines (**Fig. S1H**). CAR-NKTs expressing iC9.IL18 and iC9.IL15.IL18 mediated significantly higher levels of cytotoxicity than iC9.Q8 and/or iC9.IL15 control cells at 1:2 and 1:4 effector-to-target (E:T) ratios in both cell lines (**Fig. 1F**). These results show that NKT cells maintain a high level of IL-18Rα relative to other immune cells, with expression levels remaining unchanged between one and two stimulations, and that expressing IL-18 in GD2.CAR-NKTs provides a greater boost to *in vitro* cytotoxicity against NB than IL-15.

### IL-18 expression enhances GD2.CAR-NKT proliferation, antitumor activity, and cytokine secretion in a repeat tumor challenge assay

To determine how IL-18 and IL-15/IL-18 co-expression impacts GD2.CAR-NKT cytotoxicity after multiple encounters with tumor cells, we employed an *in vitro* repeat tumor challenge assay in which CAR-NKTs are serially stimulated with NB tumors every three-to-four days (**Fig. S2A**). CAR-NKTs expressing iC9.IL18 and iC9.IL15.IL18 mediated significantly higher levels of cytotoxicity than iC9.Q8 and iC9.IL15 controls at multiple E:Ts after three and five challenge cycles with CHLA255 and CHLA136 lines (**Fig. 2A,C**). Additionally, CAR-NKTs expressing these constructs proliferated significantly more than controls as measured after five challenge cycles and cumulatively over the course of the assay in a representative donor (**Fig. 2B, D; Fig. S2B, C**). Next, we stained for apoptosis marker annexin V after one, three, and five challenge cycles and did not detect significant differences among groups, suggesting that iC9.IL18 and iC9.IL15.IL18 do not increase CAR-NKT numbers by decreasing apoptosis rate (**Fig. S2D**). Evaluating the cytokine secretion profile of CAR-NKTs after one and three challenge cycles revealed significant increases in IFNγ, TNFα, and GM-CSF secretion from iC9.IL18 or iC9.IL15.IL18 transduced CAR-NKTs compared to iC9.Q8 and iC9.IL15 controls, while IL-2 and IL-10 were secreted at similar levels by all groups and IL-4 showed variation between groups only after one round of stimulation (**Fig. 2E**). The IFNγ-to-IL-4 ratio was similar across groups after one stimulation but after three rounds, CAR-NKTs expressing iC9.IL15, iC9.IL18, and iC9.IL15.IL18 had significantly higher ratios than the iC9.Q8 control, suggesting a Th1 skew in the former groups (**Fig. S2E**). These results demonstrate that IL-18 co-expression alone and with IL-15 enhances GD2.CAR-NKT proliferation, cytotoxicity, and production of inflammatory cytokines more than IL-15 alone after serial tumor challenge.

**Figure 2.**
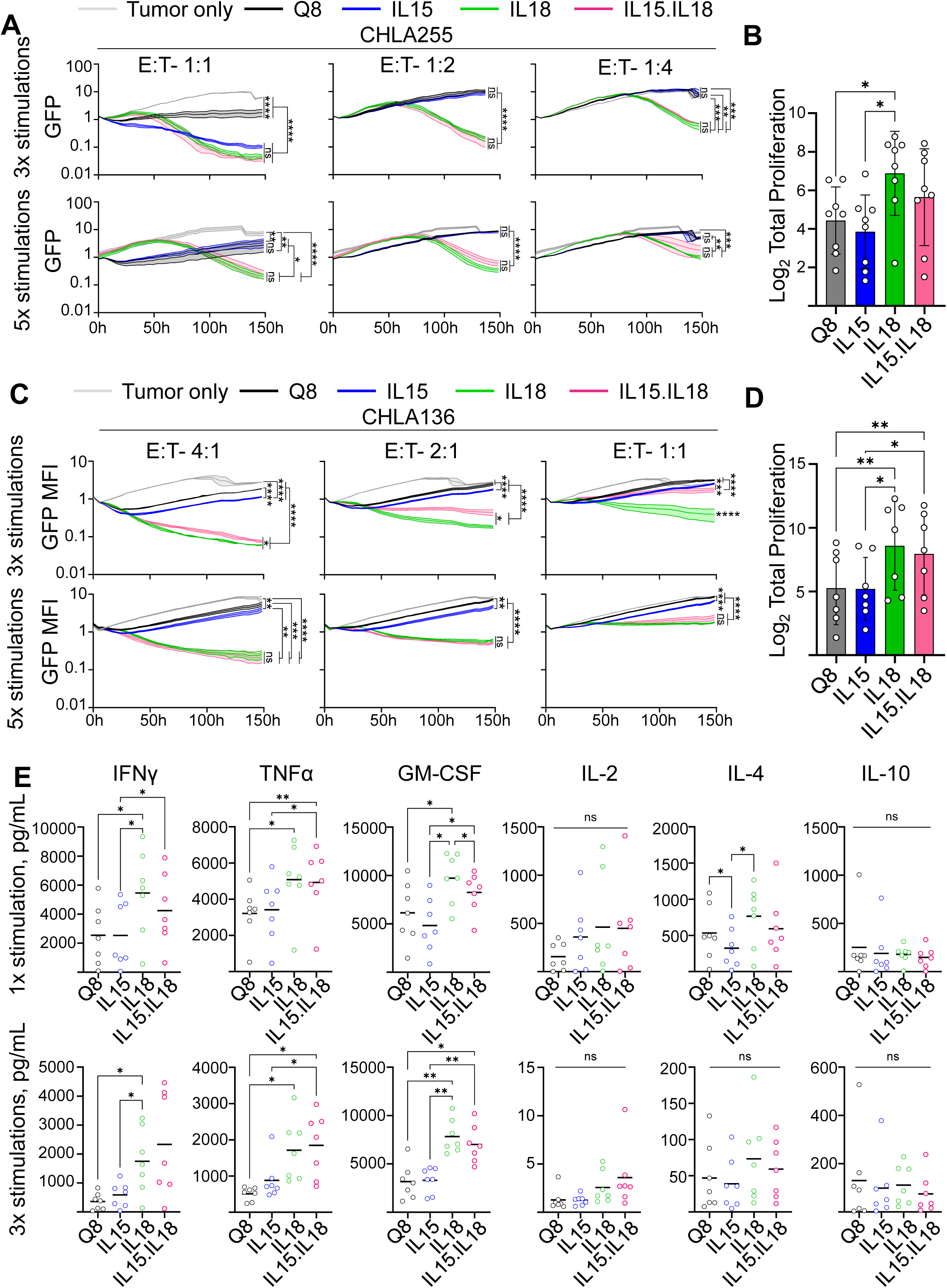
iC9.IL18 and iC9.IL15.IL18 enhance GD2.CAR-NKT *in vitro* antitumor cytotoxicity and proliferation after repeated tumor stimulation. **A**. After three (top) and five (bottom) cycles of *in vitro* tumor stimulation, GD2.CAR-NKT cells were rested overnight in fresh media then co-cultured with CHLA255-GFP cells at the indicated E:T ratios. To determine tumor cytotoxicity, changes in GFP fluorescence were measured over the indicated period using the Incucyte system. Shown are results from one representative donor of four in two independent experiments with technical triplicates for each condition. Mean (SD), ns=not significant, * *p* < 0.05, ** *p* < 0.01, *** *p* < 0.001, **** *p* < 0.0001, one-way ANOVA. **B.** Summary of NKT proliferation for indicated groups after five rounds of stimulation with CHLA255 NB cells. Mean (SD), *n*=8 donors in three independent experiments, * *p* < 0.05, paired one-way ANOVA. **C.** After three (top) and five (bottom) cycles of *in vitro* tumor stimulation, GD2.CAR-NKTs were rested overnight in fresh media then co-cultured with CHLA136-GFP cells at the indicated E:T ratios. To determine tumor cytotoxicity, changes in GFP fluorescence were measured over the indicated period using the Incucyte system. Shown are results from one representative donor of four in two independent experiments with technical triplicates for each condition. Mean (SD), ns=not significant, * *p* < 0.05, ** *p* < 0.01, *** *p* < 0.001, **** *p* < 0.0001, one-way ANOVA. **D.** Summary of NKT proliferation for indicated groups after five rounds of tumor stimulation. Mean (SD), *n*=7 donors in three independent experiments, * *p* < 0.05, ** *p* < 0.01, paired one-way ANOVA. **E.** Cytokine production after one (top) or three stimulations (bottom) with CHLA255 NB cells. Mean is shown, *n=*7 donors in three independent experiments. ns=not significant, * *p* < 0.05, ** *p* < 0.01, paired one-way ANOVA.

### IL-18 expression promotes superior GD2.CAR-NKT antitumor activity without toxicity

Next, we sought to determine whether IL-18 co-expression improves GD2.CAR-NKT antitumor activity in an *in vivo* xenogeneic mouse model of NB. NSG mice were intravenously injected with 1×10^6^ firefly luciferase-positive CHLA255 cells followed seven days later with 2×10^6^ GD2.CAR^+^ NKTs co-transduced with iC9.IL18 or iC9.IL15.IL18 with iC9.Q8, iC9.IL15, and non-transduced NKTs as controls (**Fig. 3A**). We found that both iC9.IL18 and iC9.IL15.IL18 increased the antitumor activity of GD2.CAR-NKTs compared to iC9.Q8 and iC9.IL15-treated groups based on tumor growth (**Fig. 3B,C**). However, mice treated with iC9.IL15.IL18 cells experienced weight loss 12 days after CAR-NKT infusion and had to be treated with rimiducid (CID) or vehicle control, allowing for a built-in assessment of iC9 safety switch efficacy (**Fig. 3D**). CID-treated mice rapidly regained weight, while those that received vehicle control continued to lose weight and had to be sacrificed. Flow cytometry analyses revealed a significantly higher number of human (h)CD45^+^ cells in the spleens and peripheral blood of iC9.IL15.IL18 mice (treated with vehicle control) compared to iC9.Q8 mice (**Fig. S3A,B**). In iC9.IL15.IL18 mice, CID treatment significantly reduced hCD45^+^ and Q8^+^ cell numbers compared to counts in vehicle control mice (**Fig. S3B,C**). Mice treated with CID also showed lower levels of serum cytokines including IL-15, IL-18, IFNγ, TNFα, mouse IL-6, and chemokines MIP-1α and MIP-1β compared to vehicle control mice, further indicating depletion of iC9-expressing cells (**Fig. S3D**). In mice treated with iC9.IL18 we observed significantly increased serum levels of IL-18 and IFNγ compared to iC9.Q8 or iC9.IL15 groups. Lastly, mice in the iC9.IL18 and CID-treated iC9.IL15.IL18 groups survived significantly longer than all other groups, however the vehicle control-treated iC9.IL15.IL18 group did not survive as long as the control iC9.Q8 group (*p=*0.002) (**Fig. 3E**). Based on these results, we excluded the iC9.IL15.IL18 construct from further testing and focused on evaluating iC9.IL18.

**Figure 3.**
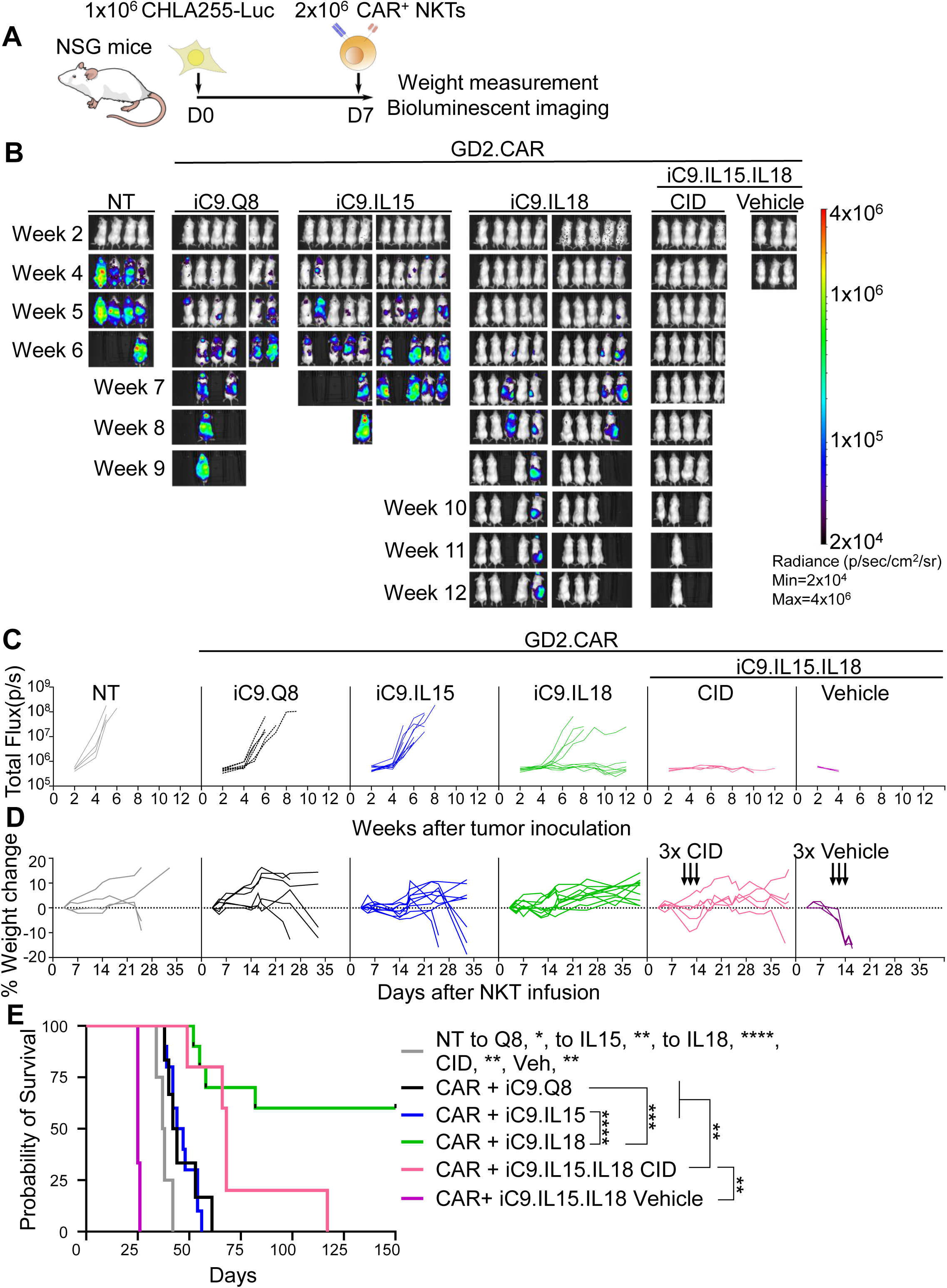
iC9.IL18 endows CAR-NKT cells with superior antitumor activity in a mouse xenograft model of NB without toxicities. **A.** NSG mice were injected via tail vein with luciferase-expressing CHLA255 NB cells, and CAR-NKTs were infused through the same route seven days later. Tumor growth was assessed weekly by bioluminescence imaging. Weight was measured every other day. **B.** CHLA255 tumor progression was measured by bioluminescence imaging after administration of the indicated CAR-NKTs (*n*=3-10 mice per group) **C.** Quantification of bioluminescence signal in (**B**). **D.** Percent weight change for mice in (**B**) after NKT infusion. Timing of CID/vehicle control administration is indicated by arrows. **E.** Kaplan-Meier survival curve for mice in (**B**). * *p* < 0.05, ** *p* < 0.01, *** *p* < 0.001, **** *p* < 0.0001, Log-rank (Mantel-Cox). Shown are results from one of two independent experiments.

Given that GD2 expression levels can be heterogeneous in many NB tumors,^40^ we evaluated the *in vivo* antitumor activity of iC9.IL18 GD2.CAR-NKTs in a second mouse model of NB based on CHLA136 NB cells, which express intermediate levels of GD2 (**Fig. S4A, Fig. S1H**). Mice treated with iC9.IL18 cells had lower tumor bioluminescence signal than iC9.Q8- and iC9.IL15 mice over seven experimental weeks and did not experience weight loss (**Fig. S4B-D**). As in the CHLA255 model, mice treated with iC9.IL18 cells survived significantly longer than all other groups (**Fig. S4E**). Thus, our results show that IL-18 safely enhances GD2.CAR-NKT antitumor activity more than IL-15 in two *in vivo* NB mouse models with different levels of GD2 expression.

### IL-18 expression drives transcription of activation-related, mitochondrial, and metabolic genes in CAR-NKTs

To determine how IL-18 impacts molecular pathways in GD2.CAR-NKT cells, we performed bulk RNA sequencing of samples at baseline and after one or three stimulations with CHLA255 NB cells using IL-15-expressing CAR-NKTs as control (**Fig S5A**). A comparison of IL-18-versus IL-15-expressing CAR-NKTs revealed 2,751 differentially expressed genes (DEGs) at baseline, 2,009 after one tumor stimulation, and 1,165 after three stimulations with an adjusted p-value under 0.05 (**Fig. S5B-D**). At baseline and after one tumor stimulation, we observed significant downregulation of *TCF7* in IL-18-expressing cells compared to IL-15, suggesting an early transition from a memory-like state towards an effector state (**Fig. 4A-C**). Accordingly, we also observed downregulation of *IL7R* and *CD27,* which we have previously reported to be upregulated in memory-like NKT cells,^41,42^ in the IL-18 group after one tumor stimulation. At baseline, IL-18 expressing CAR-NKT cells showed increased expression of genes associated with effector differentiation including *BATF3*, *BATF, CD38,* and *IFNG,* and at all three timepoints, significant upregulation of activation-related genes including *IL-2RA* and *CD86* was observed (confirmed by flow cytometry at baseline)(**Fig. S5E**). In support of these observations, gene set enrichment analysis (GSEA) revealed significant enrichment of activation- and cytokine-related terms in IL-18-versus IL-15-expressing CAR-NKTs (**Fig. 4D**). Mitochondrial- and metabolism-related terms were also significantly enriched in the IL-18 group at all timepoints, and we detected signatures of MYC targets and mTOR signaling, known regulators of lymphocyte metabolism (**Figs. 4E, Fig. S5F**).^43,44^ These findings suggest that the transcriptional program driven by IL-18 in CAR-NKTs is distinct from that of IL-15, and that many of the changes associated with IL-18 expression are related to increases in CAR-NKT mitochondrial and metabolic activity at baseline and after tumor stimulation.

**Figure 4.**
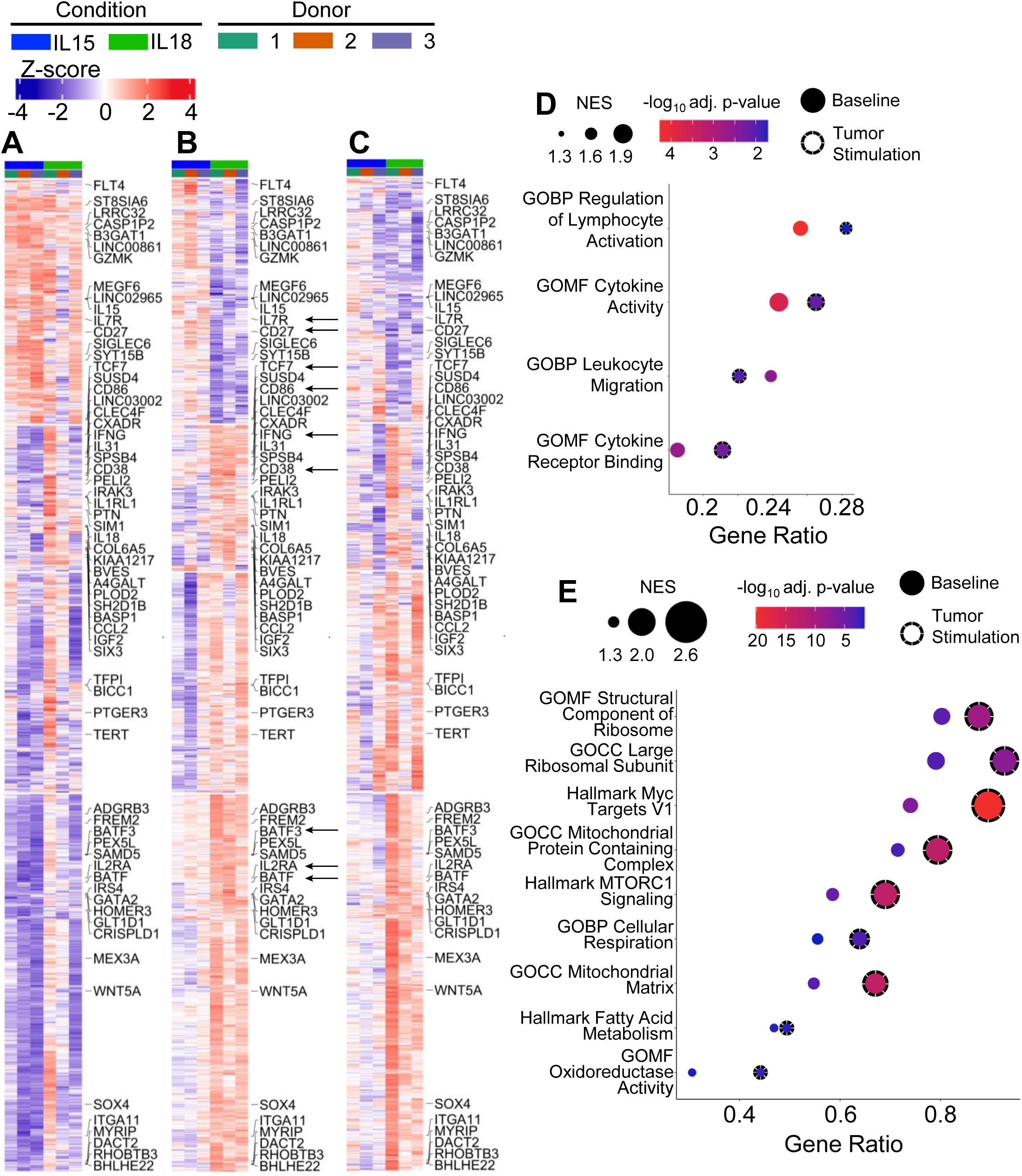
IL-18 expression drives transcription of activation-related, mitochondrial, and metabolic genes in CAR-NKTs. Gene expression heatmaps generated from CAR-NKTs expressing iC9.IL18 or iC9.IL15 at baseline (**A**) and after one (**B**) or three (**C**) tumor stimulations. Heatmaps were generated from k-means clustering of differentially expressed genes with a *p*<0.05 after one tumor stimulation. Arrows indicate genes highlighted in the body of the text. **D.** GSEA terms significantly enriched in iC9.IL18 versus iC9.IL15 CAR-NKTs at baseline (no border) and after one tumor stimulation (border). **E.** GSEA metabolism-related terms significantly enriched in iC9.IL18 versus iC9.IL15 CAR-NKTs at baseline (no border) and after one tumor stimulation (dashed border).

### Single-cell RNA sequencing identifies IL-18 specific gene clusters

We performed single cell transcriptomic analyses (scRNAseq) to define the effects of IL-18 expression on CAR-NKT transcription at single cell resolution. Using tumor-stimulated iC9.IL15 and iC9.IL18 CAR-NKT cells from three healthy donors (**Fig. S5A**), we captured a total of 21,436 cells, 58.35% of which aligned to the GD2.CAR sequence. Unsupervised single cell clustering analysis revealed the presence of eight unique expression clusters in CAR^+^ NKT cells; IL-15-expressing CAR-NKT cells were overrepresented in cluster 3, while IL-18 expressing cells were overrepresented in clusters 0, 2, and 4 (**Fig. 5A,B; Fig. S6A**). Genes upregulated in the IL-15 cluster included *DUSP2*, *BAX,* and *CISH,* a negative regulator of IL-15 signaling,^45,46^ while IL-18 clusters included *SOX4* and *MYC* (**Fig. 5C**). Next, cell cycle analysis demonstrated that cells in S phase overlapped with IL-18 clusters, consistent with the increased proliferation observed in these cells after repeated *in vitro* tumor stimulation (**Fig. 5D**). Importantly, we found that cells in IL-18-related clusters had lower expression of exhaustion gene signatures than cells in the IL-15 cluster, consistent with the observation that IL-18 drives antitumor activity, proliferation, and cytokine secretion after multiple rounds of *in vitro* tumor challenge (**Fig. 5E**).

**Figure 5.**
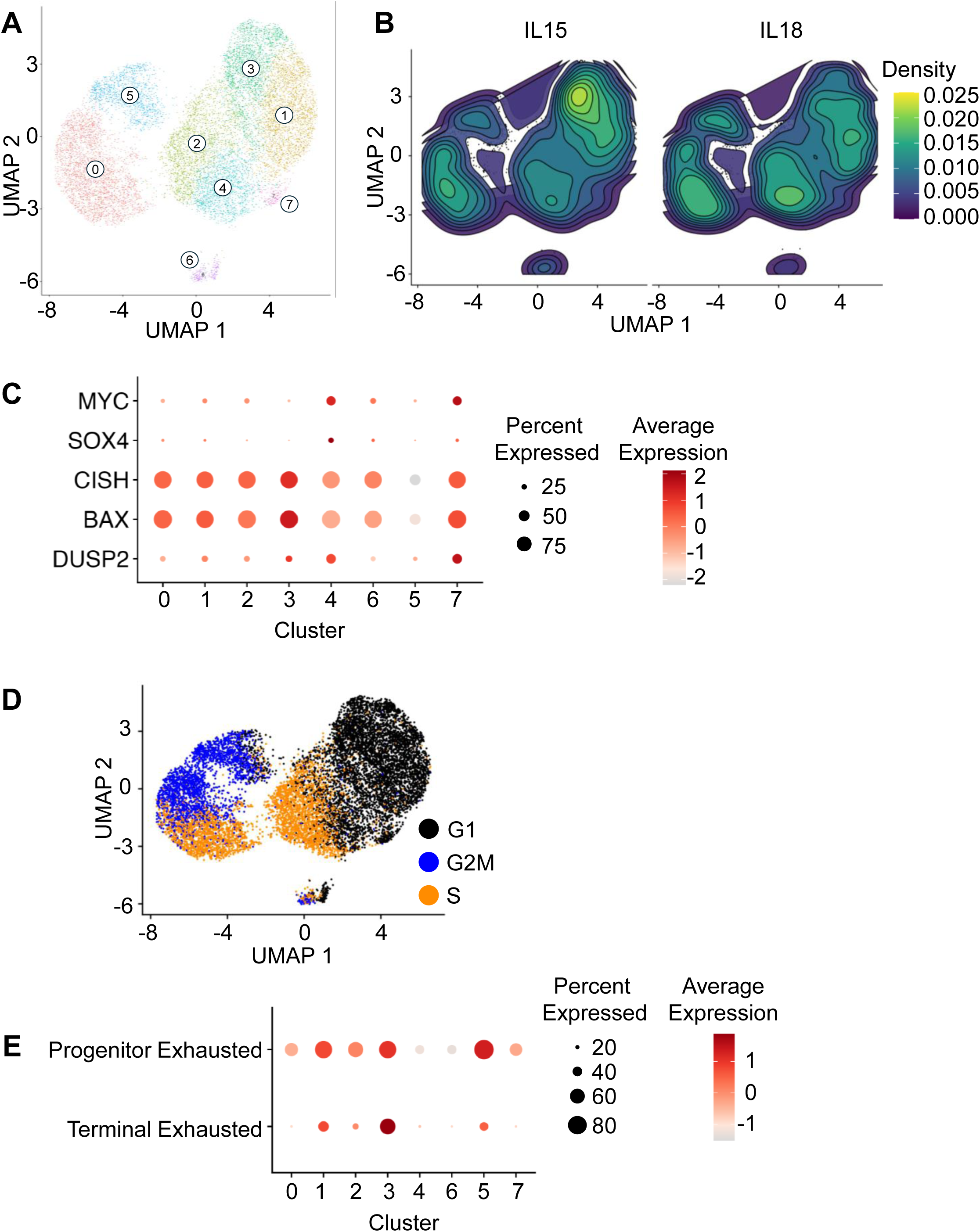
Single-cell gene expression analyses identify IL-18- and IL-15-specific gene clusters. **A.** UMAP analysis of integrated iC9.IL15 and iC9.IL18 CAR-NKT cells after one tumor stimulation **B.** Integrated data displaying density of iC9.IL15 and iC9.IL18 expressing CAR-NKT cells. **C**. Dot plot depicting differences in percent and average expression for indicated genes in clusters 1-7. **D.** Cell cycle analysis of scRNAseq data in (**A**). **E.** Progenitor and terminal exhaustion gene signature percent and average expression for clusters 1-7.

### IL-18 expression promotes oxidative phosphorylation and glycolysis in CAR-NKTs

Since our transcriptomic analyses showed enrichment of mitochondria- and metabolism-related GSEA terms in the IL-18 group, we performed several metabolic tests to further explore these findings. First, we assessed mitochondrial fitness, a key predictor of T cell function,^47^ by measuring mitochondrial mass (Mitotracker Green, MG) and membrane potential (tetramethylrhodamine ethyl ester perchlorate, TMRE) in CAR-NKTs before and after tumor stimulation. Both measures were significantly greater in CAR-NKTs expressing iC9.IL18 versus iC9.IL15 at both timepoints, and the TMRE/MG ratio was also significantly greater in iC9.IL18 CAR-NKTs, reflecting increased activity per mitochondrion (**Fig. S7A,B; Fig. 6A,B**).^48^ Given that the fatty acid metabolism term was also significantly enriched in iC9.IL18 expressing CAR-NKTs, we measured fatty acid uptake before and after tumor stimulation and found that iC9.IL18 CAR-NKTs had a significantly higher level of BODIPY uptake than iC9.IL15 cells at both time points (**Fig. S7C**). Additional targeted metabolomics analyses identified 17 differentially expressed metabolites (DEMs) in unstimulated CAR-NKTs expressing iC9.IL18 versus iC9.IL15 (*p*<0.05), which increased to 21 after stimulation (**Fig. S7D, Fig. 6C**). Stimulated CAR-NKTs expressing IL-18 showed significant enrichment of TCA cycle metabolites succinate and cis-aconitate as well as increased levels of genes encoding TCA cycle enzymes *SDHB*, *MDH2*, *MDH1*, *FH*, and *SDHA*, indicating increased TCA cycle activity (**Fig. 6D**). The glycolytic pathway also appeared to be more active in stimulated CAR-NKTs expressing IL-18 versus IL-15 based on higher levels of glycolytic intermediates phosphoenolpyruvate and glyceraldehyde 3-phosphate, significant upregulation of genes encoding glycolytic enzymes (*HK2*, *GPI*, *PFKM*, *TPI1*, *GAPDH*, *PGK1*, *PGAM1*, *ENO1, DLAT*), and higher levels of glucose uptake (**Fig. 6E, Fig. S7E**). These transcriptomic changes resulted in significant enrichment of oxidative phosphorylation and glycolysis gene signatures in IL-18 expressing CAR-NKTs before (**Fig. S7F**) and after stimulation (**Fig. 6F**); expression of these gene signatures was concentrated in the IL-18-associated clusters (0,2,4) identified in our scRNAseq analyses versus the IL-15-associated cluster (3)(**Fig. S7G**).

**Figure 6.**
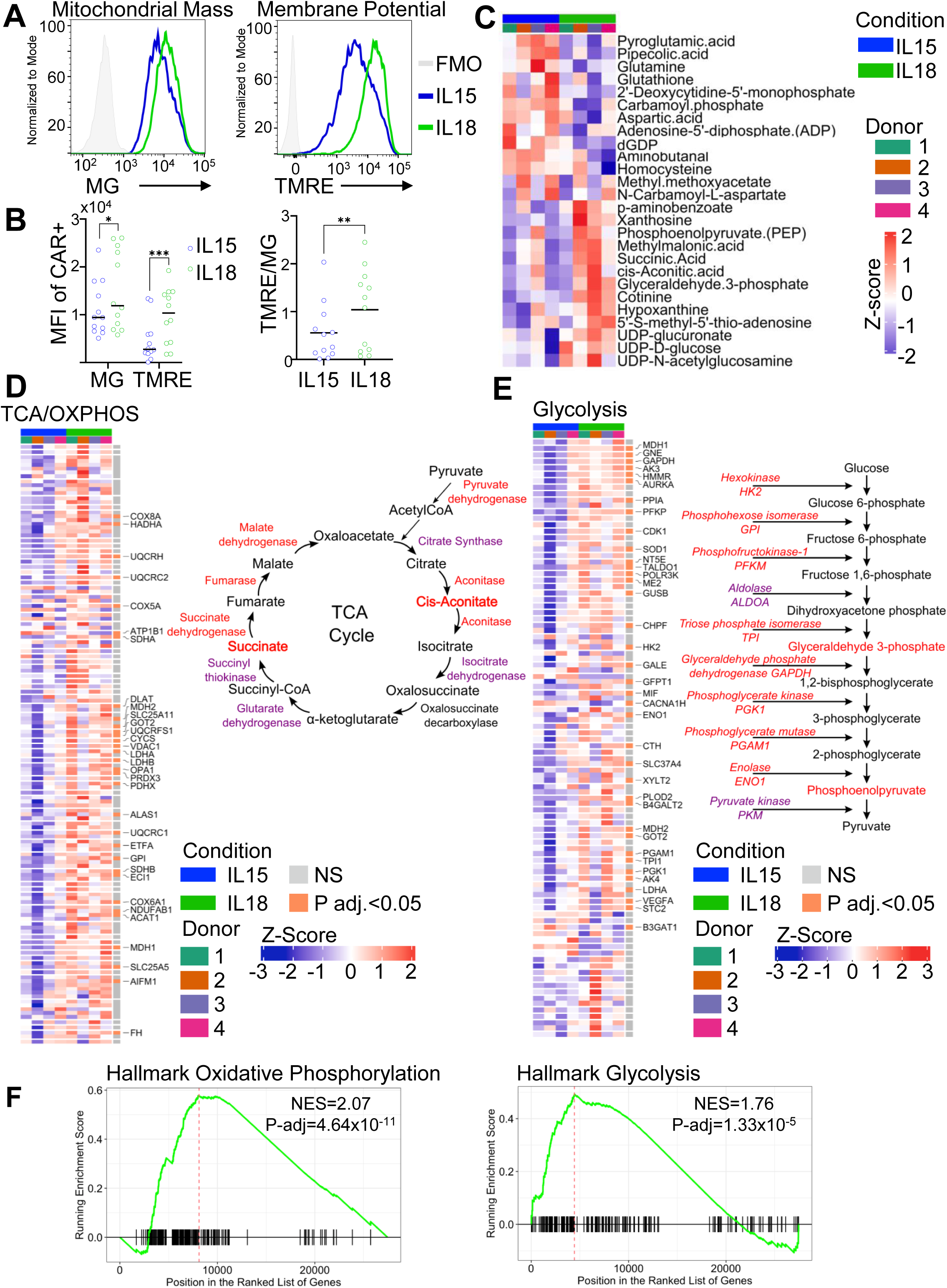
IL-18 expression broadly reprograms CAR-NKT metabolism. **A.** Flow cytometry histograms depicting MitoTracker Green (MG, left) and tetramethylrhodamine ethyl ester perchlorate (TMRE, right) staining in CAR^+^ NKT cells from a representative donor 3 days after tumor stimulation. **B.** Summary data for MG and TMRE staining after tumor stimulation (left; mean, *n*=12 donors, three independent experiments) and TMRE-to-MG ratio calculated from values in left panel (right; mean, *n*=12 donors, three independent experiments), * *p* < 0.05, ** *p* < 0.01, **** *p* < 0.0001, paired two-tailed t-test. **C.** Differentially expressed metabolites in iC9.IL18 versus iC9.IL15 expressing CAR-NKTs 24 hours after stimulation with plate-bound 1A7 antibody (*n*=4 donors, *p*<0.05). **D.** Heatmap depicting oxidative phosphorylation metabolite related genes in CAR-NKTs after one stimulation (left). TCA cycle schematic showing significantly upregulated molecules in red text. Purple italic text indicates upregulated transcripts with *p*<0.05 and adjusted *p*>0.05 (right). **E.** Heatmap depicting glycolysis metabolite-related genes in CAR-NKTs after one stimulation (left). Glycolysis pathway schematic showing significantly upregulated molecules in red text. Purple italic text indicates upregulated transcripts with *p*<0.05 and adjusted *p*>0.05 (right). **F.** Gene set enrichment analysis of oxidative phosphorylation (left) and glycolysis signatures (right) in stimulated CAR-NKTs expressing iC9.IL18 versus iC9.IL15.

To identify other metabolic pathways that are impacted by IL-18 expression, we performed pathway enrichment analysis using the DEMs we identified above and found that the *Glutamate metabolism* term was significantly enriched in iC9.IL18 CAR-NKTs before and after stimulation (**Figs. S8A,B**). Indeed, glutamine was the most downregulated metabolite in iC9.IL18 CAR-NKTs in both conditions (**Fig. S8C**) despite increased expression of glutamine transporters *SLC7A5* and *SLC1A5* (**Fig. S8D**). We also noted differential expression of transcripts related to glutamine metabolism including *GLUL*, *PPAT*, *GFPT1*, *CAD*, *GPT2,* and *GOT2.* These findings indicate activation of the glutaminolysis pathway and are consistent with the role of this amino acid in T cell activation, proliferation, and cytokine production.^43,49^ The *Purine metabolism* term was also significantly enriched in iC9.IL18 CAR-NKTs, as evidenced by differential expression of inosine, hypoxanthine, L-aspartic acid, xanthosine, glutamine, dGDP, and ADP either before (**Fig. S7D**) or after stimulation (**Fig. 6C**). Notably, inosine, which has been associated with promoting T cell fitness,^50,51^ was upregulated in iC9.IL18 CAR-NKTs before stimulation. Hypoxanthine, a byproduct of inosine metabolism, was significantly upregulated after stimulation, suggesting that this pathway remains active in both conditions. Transcriptomic analyses supported the involvement of this pathway by revealing significant increases in the level of expression for several genes involved in purine metabolism in IL-18-expressing CAR-NKTs before and after stimulation (**Fig. S8E**). Enrichment of the purine metabolism pathway is consistent with the requirement for an increase in the nucleotide pool post-T cell activation.^52,53^

Overall, our findings demonstrate that IL-18 expression has broad effects on CAR-NKT cell metabolism, including promoting oxidative phosphorylation and glycolysis, providing a mechanistic explanation for the observed increase of CAR-NKT functional fitness and antitumor activity in NB models.

## Discussion

Solid tumors remain largely resistant to therapeutic immune cell products such as CAR-redirected T, NK, and NKT cells due in part to the highly suppressive TME. A promising strategy for overcoming TME-mediated suppression is to engineer antitumor effector cells to express stimulatory cytokines.^21,23,54^ Given that cytokine receptor expression levels vary among different lymphocyte subsets and can also depend on differentiation stage and activation status,^37,55,56^ it is crucial to design cell type-specific studies for evaluating candidate cytokines that take these factors into consideration. In this study, we explored for the first time how IL-18 impacts CAR-NKT cell functionality, antitumor activity, and toxicity in preclinical NB models. Compared to IL-15, the only cytokine that has so far been evaluated clinically in therapeutic CAR-NKTs,^54^ IL-18 enhanced CAR-NKT effector function and antitumor activity in NB models while maintaining a favorable safety profile.

Surprisingly, we discovered that human NKTs expressed significantly higher levels of IL-18Rα than NK and conventional T cells, suggesting that IL-18 signaling is particularly relevant for this cell type. In both short-term cytotoxicity and serial tumor challenge assays, CAR-NKTs expressing iC9.IL18 and iC9.IL15.IL18 mediated higher levels of cytotoxicity against both GD2^high^ and GD2^int^ NB cells than iC9.Q8 and iC9.IL15 control cells. Cytotoxicity against GD2^int^ cells is of particular clinical relevance as GD2 expression is variable in NB patients, and lower GD2 expression has been observed as an escape mechanism in NB patients treated with GD2-specific monoclonal antibodies,^57^ potentially playing a role in resistance to GD2.CAR-redirected immunotherapy.

Consistent with these observations, IL-18-containing constructs also promoted robust CAR-NKT proliferation and induction of multi-cytokine responses following *in vitro* serial tumor challenge compared to iC9.Q8 or iC9.IL15. *In vivo*, IL-18-expressing CAR-NKT groups mediated superior tumor clearance compared to controls, but mice treated with iC9.IL15.IL18 experienced toxicities including increased systemic expansion of CAR-NKTs and elevated serum concentrations of human cytokines. Synergy between IL-15 and IL-18 has previously been reported in NK cells, and the NKT hyperactivation that we observed is therefore not without precedent.^58,59^ Hence, we excluded the IL-15/IL-18 combination from further consideration as a clinical candidate, focusing instead on iC9.IL18, which mediated potent antitumor activity in GD2^high^ and GD2^int^ NB models without causing evident toxicity. These findings align with recent clinical data demonstrating that IL-18-expressing CD19-CAR T cells are safe in lymphoma patients,^60^ reinforcing the feasibility of incorporating IL-18 into CAR-based immunotherapies.

Though CAR-NKTs expressing iC9.IL18 did not generate toxicity in mice, incorporating the iC9 safety switch into the final clinical construct is critical to ensuring that the therapeutic cells can be eliminated after infusion if needed. iC9 has been validated in cell therapy trials as an effective safety switch, including a recent report in GPC3-positive solid tumor patients treated with GPC3.CAR T cells co-transduced with an iC9.IL-15 construct.^54^ This study also demonstrated the advantage of a system where the CAR is encoded by one construct and the iC9 switch and cytokine by a second construct; elimination of IL-15-producing cells by iC9 activation effectively controlled toxicity while providing evidence of objective tumor control.^54^ Our results also validated the functionality and robustness of the iC9 switch, which was activated successfully following CID treatment *in vitro* and *in vivo* in mice receiving iC9.IL15.IL18 CAR-NKTs.

Importantly, bulk transcriptomics, single-cell RNA sequencing, and metabolomic profiling revealed that IL-18 induces a distinct transcriptional and metabolic program in CAR-NKTs. Compared to IL-15, IL-18 expression increased mitochondrial activity, oxidative phosphorylation, glycolysis, and glutaminolysis. These processes have been previously been linked with T cell activation^61,62^ and are likely to be important for sustained effector function of CAR-redirected immune cells at tumor sites. Indeed, the ability of IL-18 expressing CAR-NKTs to exploit different metabolic pathways and nutrients could prove advantageous for persistence in the nutrient-poor TME, and evaluating the roles of specific metabolites in CAR-NKT function would provide a focus for future studies.

In conclusion, our findings demonstrate that IL-18 enhances the antitumor efficacy of CAR-NKT cells by promoting metabolic fitness and effector function without increasing off-target toxicity. These results support the clinical translation of IL-18-expressing CAR-NKT cells for the treatment of NB and potentially other solid tumors. Future studies should focus on optimizing IL-18-based CAR-NKT therapies to maximize their therapeutic potential while minimizing toxicity, paving the way for more effective and durable immunotherapies for solid tumors.

## Methods

### Cell lines

CHLA255 and CHLA136 NB cell lines were cultured in IMDM medium supplemented with 20% FBS and GlutaMAX (Gibco, 35050061). HEK293T cells were maintained in IMDM supplemented with 10% FBS and GlutaMAX. Luciferase-transduced or GFP-transduced CHLA255 or CHLA136 cell lines were used in indicated experiments. K562 artificial antigen presenting cells (aAPCs)^42^ were maintained in RPMI1640 supplemented with 10% FBS and GlutaMAX. All cell lines were routinely tested for Mycoplasma contamination. Cell line identities were validated at MD Anderson Cancer Center (Houston, TX) using the short tandem repeat method within one year of use.

### Plasmids

The GD2.28ζ plasmid has been previously described.^63^ iC9, cytokines, and Q8 sequences were cloned into the SFG gamma retroviral backbone^64^ using NEBuilder® HiFi DNA Assembly master mix (NEB, E2621L). The IL-2-SP-IL-18 sequence was synthesized by IDT. iC9,^65^ IL-15,^22^ and Q8^36^ sequences were PCR-amplified from existing constructs using Platinum™ SuperFi II PCR master mix (ThermoFisher, 12368010). The pNFKB reporter construct was generated by PCR amplifying from the NFKB-eGFP from pSIRV-NF-kB-eGFP construct (Addgene: 118093)(gift from Peter Steinberger),^66^ and inserting the fragment into a lentiviral backbone by HiFi DNA assembly, followed by an SFFV promoter and the tEGFR transduction marker. The psPAX2 and pMD2.G plasmids (Addgene:12260 and 12259, respectively) were gifts from Didier Trono.

### Virus production

Retroviral particles were generated as previously described.^67^ Lentiviral particles were generated as previously described^68^.

### NKT manufacturing and transduction

NKTs were isolated from the buffy coats of healthy human donors (Gulf Coast Regional Blood Center) and expanded as previously described.^37^ Six days after isolation, NKTs were retrovirally transduced with iC9/cytokine constructs and the following day, iC9-NKTs were collected and transduced with the CAR construct. For pNFKB reporter construct experiments, equal volumes of CAR (retroviral) and reporter (lentiviral) supernatants were mixed and spun down at 4000xg for 1.5 hours at 32C onto retronectin-coated non-tissue culture-treated 24-well plates. Two days after the second transduction, NKT cells were transferred to 24-well G-Rex plates and stimulated with irradiated K562 aAPCs^42^ in the presence of 100 ng/mL α-galactosylceramide (αGalCer, Avanti Polar Lipids, 86700P-1mg) and 200U/mL IL-2 (National Cancer Institute/TECIN™, 23-6019). NKT cells were expanded for an additional 10 to 12 days and then used immediately or cryopreserved for subsequent experiments.

### *In vitro* cytotoxicity

*In vitro* cytotoxicity of NKT cells co-transduced with the CAR and IL-18 and/or IL-15 was evaluated using an Incucyte S3 imaging system. One day before starting the assay, 1×10^4^ GFP-expressing NB cells were plated in each well of a flat-bottom tissue culture-treated 96-well plate and incubated for five minutes at room temperature to allow for even distribution of cells on the bottom of the wells. On the same day, CAR-transduced and non-transduced NKT cells were thawed and recovered overnight in medium containing 200u/mL of IL-2. The next day, NKT cells were washed and resuspended in fresh medium without cytokines. NKT cells were serially diluted to achieve the indicated effector-to-target ratios and added to the NB cells plated the day before. Incucyte imaging was performed every six hours for seven days. Images were analyzed using Incucyte software (Sartorius).

### Repeat tumor stimulation assay

Cryopreserved NKTs were thawed and rested overnight in the presence of 200u/mL IL-2. The following day, cells were washed, resuspended in fresh medium containing 50u/mL IL-2, and 3×10^6^ CAR^+^ NKT cells were plated with an equal number of tumor cells per well of a 24-well G-Rex. Total live CAR^+^ NKT cell and tumor cell numbers were determined every three-to-four days (CHLA255 model) or four-to-five days (CHLA136 model) by flow cytometry using CountBright Plus counting beads (ThermoFisher, C36995). After counting, 3×10^6^ CAR^+^ NKT cells were replated with an equal number of tumor cells for a total of five tumor stimulation rounds. After one or three tumor stimulations, NKT cells were rested overnight in fresh medium containing 200u/mL IL-2 for *in vitro* cytotoxicity testing at different ratios with NB cell lines or for RNA isolation. NKT cells were rested for two additional days for flow cytometry phenotyping. Evaluation of cytotoxicity after repeat tumor stimulation was performed as described in the *In vitro cytotoxicity* section.

### Luminex assays

NKT cells were washed and resuspended in fresh medium without IL-2 and cultured in 96-well plates with or without 1×10^5^ NB cells at a 1:1 E:T and at a final concentration of 0.5×10^6^ CAR^+^ NKT cells/mL. Supernatants were collected after 24 (9-plex) or 48 (2-plex) hours and stored at −80C until analysis. Supernatants were analyzed using a 2-plex (IL-15 and IL-18) or 9-plex (IL-15, IL-18, IFNγ, TNFα, IL-4, IL-6, IL-10 and IL-2) MILLIPLEX® Human Cytokine/Chemokine/Growth Factor Panel A magnetic bead panel (HCYTA-60K, Millipore) according to the manufacturer’s instructions. Serum samples from mice were collected on day 20 after NKT infusion and stored at −80C until \ analysis. Samples were analyzed using a 1-plex (mouse IL-6) MILLIPLEX® Mouse Cytokine/Chemokine Magnetic Bead Panel (MCYTOMAG-70K-1, Millipore) and a 9-plex (human IL-18, IL-15, IFNγ, GM-CSF, G-CSF, IL-4, TNFα, MIP-1α, and MIP-1β) MILLIPLEX® Human Cytokine/Chemokine/Growth Factor Panel A (HCYTA-60K-10, Millipore). To facilitate analysis of samples that had undetectable cytokine/chemokine levels, those samples were assigned a value of half the detection limit specified by the respective kit. All Milliplex assays were analyzed using the Luminex FLEXMAP 3D system with xPONENT software (Luminex) according to the manufacturer’s instructions.

### Flow cytometry

Spleen samples were homogenized using a gentleMACS dissociator and tubes (Miltenyi Biotech). Blood samples were collected via tail vein. Spleen and blood samples were treated with ACK lysis buffer for five minutes at room temperature and then washed twice in DPBS. Processed tissue samples or cultured cells were stained with fixable viability dye eFluor 780 (eBioscience, 65-0865-18). NKT phenotype was assessed using the following antibodies: AF488 hCD45 (HI30), PE-Cy7 or BV421 iNKT (6B11), PE-Cy7 CD3 (UCHT1), BV785 CD39 (A1), PE IL-18Rα (H44), PeCy7 CD25 (M-A251), PE CD86 (IT2.2), BV421 Mouse IgG2a (MOPC-173, Biolegend), BV480 mouse CD45 (30-F11), BV421 CD62L (DREG-56), BV650 TIM3 (7D3), BUV395 CD4 (SK3), BV421 GD2 (14.G2a, BD), PE PD-1 (J105), BUV737 LAG3 (3DS223H), PerCP-eFluor-710 TIGIT (MBSA43, eBioscience), and PE-CD34 (QBEND/10, Invitrogen). GD2-CAR expression was evaluated using an AF647 14G2a anti-idiotype 1A7 antibody purified from an 1A7 mouse hybridoma (ATCC, HB-11786), which was custom-conjugated to Alexa Fluor 647 by BioLegend. Apoptosis was assessed by staining NKT cells with V500 Annexin V (561501, BD) in 1x apoptosis buffer (BD) supplemented with 2% FBS for 15 minutes at room temperature then washing the cells and resuspending in 1x apoptosis buffer for analysis. Analysis was performed on FACSymphony A5 (BD), FACSymphony A1, or iQue 3 (Sartorious) flow cytometers. Flow cytometry data was further analyzed using Flowjo v10.10 software (BD).

### Metabolic status flow cytometry

Briefly, NKT cells were washed in DPBS and resuspended in fresh NKT medium and cultured for 30 minutes at 37C before adding Mitostatus TMRE (564696, BD) and Mitotracker green (M7514, Invitrogen) to a final concentration of 10nM. To assess glucose uptake, NKT cells were washed and resuspended in glucose-free RPMI 1640 medium supplemented with GlutaMAX (61870036, Gibco) and incubated for 30 minutes at 37C, after which cells were stained with 2-NBDG (Cayman chemical, 11046) at a final concentration of 50 µM. To assess fatty acid uptake, NKT cells were washed in DPBS and resuspended in RPMI 1640 medium supplemented with GlutaMAX and 20 µM of fatty acid-free BSA (126575, Millipore-Sigma). The cells were then incubated for 30 minutes at 37C before staining with BODIPY™ 500/510 C_1_, C_12_ (D3823, Invitrogen) at a final concentration of 0.5 µM. After adding the respective dyes, the cells were further incubated at 37C for 30 minutes in the dark then washed twice in DPBS containing 2% FBS and stained with fixable viability dye 780 and surface antibodies. Samples were then analyzed by flow cytometry as described in the previous section.

### Phosphoflow staining

NKT cells were washed and resuspended in complete medium (without exogenous IL-2) at 0.25×10^6^ NKTs per mL and 0.5×10^6^ cells were plated per well of a 24-well plate. Five days later, the cells were collected and stained with fixable viability dye eFluor 780 at 4C for 15 minutes. Cells were then washed in cold PBS and stained for surface markers. NKTs were further processed using the Transcription Factor Phospho set (563239, BD) according to manufacturer’s instructions. In brief, cells were fixed for 1 hour at 4C in 1x Fix-Perm buffer and then washed twice in 1x Perm-Wash buffer. Samples were then resuspended in Perm Buffer III (stored at −20C) and incubated on ice for 20 minutes. Cells were then washed twice in 1x Perm-Wash buffer and then stained overnight at 4C with PE-pSTAT5 (pY694, 562077, BD) at a 1:15 dilution in 1x Perm-Wash buffer. After an overnight incubation, cells were washed twice with Perm-Wash buffer and then resuspended in MACS buffer (DPBS + 0.5% FBS + 1mM EDTA). Analysis was performed as described in the *Flow Cytometry* section.

### Bulk RNAseq and data processing

CAR^+^ NKT cells were enriched by fluorescence-activated cell sorting after manufacturing. For experiments performed after tumor challenge, first we confirmed tumor elimination by flow cytometry and then we enriched live non-apoptotic CAR-NKTs by depletion of annexin V^+^ cells using the EasySep™ Dead Cell Removal (Annexin V) Kit (STEMCELL Technologies, 17899) following the manufacturer’s instructions. Cells were washed, spun down, and lysed in TRI Reagent (R2050-1-200, Zymo Research) followed by RNA isolation using the Direct-zol RNA Mini Prep kit (R2053, Zymo Research). Library generation and sequencing were performed by Novogene using a NovaSeq X Plus (Illumina) with 150bp paired-end read length and around 50 million read-pairs per sample. Fastq reads were first trimmed of adapters using Trimmomatic v0.39 under the default settings using adapters from TruSeq2-PE.fa.^69^ Each of the 150bp paired-end RNAseq reads for each sample were aligned to the hg38 reference genome using STAR v2.7.^70^ Aligned reads were quantified using FeatureCounts,^71^ and differential gene analysis was performed for each CAR condition within each cycle, correcting for donors using DESeq2 v1.42.1 with a false discovery rate <0.05.^72^ To create heatmaps, significant differentially expressed genes (DEGs) were selected and standardized with *z*-scores across samples. Hierarchical clustering and visualization were implemented using ComplexHeatmap.^73^ Pathway enrichment was performed using clusterProfiler v4.10.0^74^ through gene set enrichment analysis (GSEA) using signature genes from the C5 ontology gene sets from MSigDB.^75^

### Single-cell RNAseq

CAR-NKTs co-transduced with IL-15 or IL-18 from three healthy donors were stimulated 1:1 with CHLA255 NB cells for three days as previously described, rested overnight in 200u/mL IL-2, and dead cells were then removed as previously described. A total of 10,000 cells per condition per donor were barcoded, and sequencing libraries were generated using the Chromium GEM-X Single Cell 5’ v3 Gene Expression kit (10x Genomics) following the manufacturer’s instructions. Libraries were sequenced by Novogene on an Illumina NovaSeq X Plus instrument with an average read depth of 30,000 reads per cell. The GD2.CAR sequence was incorporated into the GRCh38 human reference genome for data pre-processing and read alignment. Reads were aligned according to the 10X Genomics standard using Cell Ranger (v8.0.1)^76^ and downstream analyses were performed with Seurat (v5.0.2).^77^ We obtained an average of 6.13×10^7^ total reads per sample. Cells were classified as CAR positive if they could be aligned to the GD2.CAR sequence. Only cells with raw unique molecular identifier (UMI) counts >0 were used for analysis. Further gene expression and cell filtering were performed under the following exclusion criteria: features with <10 cells, cells with <200 genes, cells with >10% of mitochondrial raw UMIs, and cells with > 7,000 genes. Sample integration was performed using Harmony (v1.2.0).^78^ We performed principal component analysis on scaled data and unsupervised Louvain clustering of CAR^+^ or total cell populations from IL-18 and IL-15 CAR samples with a resolution of 0.5. Differential gene expression was performed on CAR^+^ cells to compare clusters by aggregating cells from those clusters and testing for DEGs. DEG analysis was performed by implementing FindMarkers(min.pct=0.2, min.cells.feature=100, test.use=”DESeq2”) in Seurat, and DEGs were defined as those with a BH-adjusted *p*-value <0.05. Gene expression scores and per-cluster GSEA enrichment were performed using gene sets from Hallmark or from predefined gene sets by Miller^79^ using AddModuleScore() in Seurat. Enrichment analysis was performed with clusterProfiler,^80^ and significantly enriched terms were defined as those with a false discovery rate <0.05.

### Metabolic profiling

Unstimulated cryopreserved CAR-NKT cells were thawed and rested overnight in medium containing IL-2. Cells were then counted, washed three times in cold DPBS, counted again, and 5×10^6^ cells were transferred to a microcentrifuge tube to be spun down and frozen. To avoid confounding effects from tumor-generated metabolites, we stimulated CAR-NKT cells using plate-bound 1A7 anti-idiotype monoclonal antibody. Briefly, non-tissue culture treated 24-well plates were coated with 1µg of 1A7 antibody diluted in PBS at 1 µg/mL, and plates were incubated overnight at 4C. Plates were then washed with PBS and blocked for 30 minutes at room temperature using complete NKT medium, after which medium was removed by aspiration. Next, 0.5×10^6^ NKT cells were loaded into the wells and the plates were spun down at 400xg with a low deceleration rate and placed in an incubator. One day later, stimulated NKTs were collected and processed similarly to unstimulated CAR-NKTs. Metabolites were extracted from cell pellets using a liquid-liquid extraction method as previously described.^81–83^ Pooled samples were utilized for quality control. In brief, cells were homogenized in a methanol-water (4:1) solution containing an internal standard mix. Following homogenization, chloroform and water were added to facilitate phase separation. Mass spectrometry data were collected using multiple reaction monitoring on a 6495 Triple Quadrupole mass spectrometer coupled to a high-performance liquid chromatography system (Agilent Technologies), operated with Agilent Mass Hunter Software.^81–86^ Detailed liquid chromatography-mass spectrometry conditions and parameters were consistent with prior studies.^81^ Peak integration and data analysis were performed using Agilent Mass Hunter Quantitative Analysis software. The extracted peak areas were log2-transformed and normalized using isotopically labeled internal standards Jasmonic acid, L-Tryptophan, and L-Zeatin specific to each method. For analysis, each batch of standardized metabolites was analyzed separately by explicitly defining internal batch standards using the estimateSizeFactors() function in DESeq2^72^. We tested metabolites from IL-18 versus IL-15 samples either at baseline or after stimulation time points and controlled for donor variation using DESeq(fitType=”local”) in DESeq2. We then appended the differentially expressed metabolites (DEMs) output to their respective condition to examine overall changes in metabolite expression. DEMs were identified using a *p*<0.05 for descriptive visualization purposes. Pathway enrichment analysis was performed using DEMs in the IL-18 versus IL-15 group with the Enrichment Analysis tool from MetaboAnalyst.^87^

### In vivo studies

NSG mice were obtained from The Jackson Laboratory and maintained at the Baylor College of Medicine animal care facility. Eight-to ten-week-old mice were injected via tail vein with 1×10^6^ firefly luciferase-expressing CHLA255 or CHLA136 NB cells. Seven days later, mice were intravenously infused with 2×10^6^ or 1×10^7^ CAR-NKTs. Mice received intraperitoneal (i.p.) injections of IL-2 (2000 U/mouse) every other day for two weeks. Tumor growth was assessed weekly by bioluminescent imaging using an IVIS Lumina III *in vivo* imaging system (Revvity). Toxicity was assessed by measuring animal weight every other day. Mice were treated i.p. with AP1903 dimerizer (HY-16046, Medchem Express) or vehicle if weight loss was ≥10% of pre-NKT infusion measurements. Blood samples for cytokine and flow cytometry analyses were collected after AP1903 treatment. After dimerizer or vehicle treatment, spleen samples were collected for further flow cytometry analyses. Animal experiments were performed according to IACUC-approved protocols (BCM IACUC protocol AN-5194).

### Statistics

For *in vitro* or *in vivo* experiments comparing two groups, two-tailed paired or unpaired t-tests were used. When comparing more than two groups, unpaired or paired ANOVA with Tukey’s or Dunnett’s multiple comparison tests were used. Survival curves were compared using the log-rank Mantel-Cox test. Statistical methods were not used to pre-determine sample sizes. Unless otherwise indicated, statistical analyses were calculated using GraphPad Prism v10.4.1.

## Data availability

Raw bulk and single cell RNA sequencing, and raw metabolomics data collected are available from the corresponding author upon request.

## Acknowledgments

This work was supported by the NIH National Cancer Institute grants: R01-CA262250 (L.S.M.), R01-CA247436 (G.D.); and Isabella Santos Foundation (L.S.M). The authors are grateful for the excellent technical assistance provided by the staff at the Flow Cytometry Core Laboratory of the Texas Children’s Cancer and Hematology Center, Small Animal Imaging Core facility at Texas Children’s Hospital and the Metabolomics Core at Baylor College of Medicine (CPRIT-RP210227 and NIH P30 CA125123). Illustrations were edited and generated using Inkscape software, some illustrations were obtained from the NIAID Visual & Medical Arts BIOART source.

## Author contributions

Conceptualization, G.B., G.D. and L.S.M., Methodology, G.B., E.L., G.T., X.R., X.X., A.N.C., G.D. and L.S.M., Investigation, G.B., E.L., K.D., P.H., L.D.C., Y.W., B.Y., L.G., M.W., Formal analysis, G.B. and D.A.C., Writing – original draft, G.B. and L.S.M., Writing – review & editing, G.B., A.N.C., E.J.D.P, and L.S.M., Visualization, G.B. and D.A.C.

## Declaration of interests

G.B., A.N.C, and L.S.M. are co-inventors on a pending patent application relevant to this work. Other authors declare no potential conflicts of interest.

## Supplementary Figures

**Supplementary Figure 1. Generation and characterization of NKT cells co-transduced with a GD2.CAR and an iC9.cytokine construct**

**A.** Two days after GD2.CAR transduction, co-transduced NKTs were stimulated using αGalCer loaded aAPCs and expanded for 10 days. Then, NKT purity was assessed by flow cytometry analysis using antibodies specific to the NKT TCR and CD3. Representative flow cytometry analysis of NKT purity 10 days after transduction. **B.** Quantification of CAR^+^ NKT cell expansion after 10 days of secondary stimulation. Mean is shown, *n*=15 donors in four independent experiments, * *p* < 0.05, one-way paired ANOVA. **C.** Quantification of transgenic cytokine production after tumor stimulation, *n*=6 donors in 3 independent experiments. **D.** CAR-NKT cells were cultured for three days without IL-2, and expression of phosphorylated STAT5 was evaluated by intracellular flow cytometry staining. Results from a representative donor (left), and summary data from *n*=5 donors (right). Mean is shown, * *p* < 0.05, paired one-way ANOVA. **E.** CAR and iC9.cytokine expressing NKTs were transduced with a lentivirus encoding an NF-kB reporter. After, cells were stimulated with αGalCer loaded aAPCs and expanded for 10 days. Then NF-kB reporter activation was measured by flow cytometry. Representative quantification of NF-kB activity in CAR-NKTs using a reporter construct (left), and summary data of from *n*=5 donors. Mean is shown, * *p* < 0.05, paired one-way ANOVA. **F.** CAR-NKT cells co-transduced with the indicated constructs were treated with 1nM rimiducid/AP1903 (CID) or vehicle control, and Q8 marker expression was evaluated one day later by flow cytometry. Representative flow cytometry plots of Q8/CAR expression are shown. **G.** Summary data of changes in Q8 expression from (**F**) in *n*=6 donors and two independent experiments. **** *p* < 0.0001, paired two-tailed t-test. **H.** Flow cytometry histograms of GD2 expression in the indicated NB lines or NKT cells.

**Supplementary Figure 2. iC9.IL18 expression enhances CAR-NKT function after repeated tumor challenge**

**A.** Repeat tumor stimulation assay experimental design. CAR-NKTs are co-cultured with GFP^+^ NB cells for three-to-four days, counted by flow cytometry, and replated 1:1 with NB for the next stimulation for a total of five rounds. After selected stimulation rounds, CAR-NKTs are rested overnight with IL-2 and prepared for cytotoxicity assay. **B.** Cumulative CAR^+^ NKT cell fold change in a representative donor over five rounds of stimulation with CHLA255 NB cells. **C.** Cumulative CAR^+^ NKT cell fold change in a representative donor over five rounds of stimulation with CHLA136 NB cells. **D.** After one, three, or five stimulation cycles with CHLA255 cells, CAR-NKTs were rested for three days in fresh media containing 200 u/mL IL-2 and apoptosis was assessed by annexin V staining. Mean is shown, *n=4* donors in three independent experiments, ns=not significant, paired one-way ANOVA. **E.** IFNγ-to-IL-4 ratios calculated from cytokine concentrations shown in Fig. 2E after one or three tumor stimulation rounds. Means are shown, *n=*7 donors in three independent experiments, ns=not significant, * *p* < 0.05, paired one-way ANOVA.

**Supplementary Figure 3. Characterization of iC9.IL15.IL18 toxicities in mice**

**A-D.** Three days after the last CID/Vehicle dose, mice blood, serum and spleens were collected for further characterization. **A.** Representative flow cytometry quantification of human CD45^+^ cells in the spleens of mice from Fig. 3B treated with GD2.CAR-NKT cells co-transduced with iC9.Q8 or iC9.IL15.IL18. **B.** Quantification of hCD45+ cells in the blood of mice from Fig. 3B treated with the indicated constructs, *n*=3, * *p* < 0.05, ** *p* < 0.01, ordinary one-way ANOVA. **C.** Quantification of Q8 cells in the blood of mice that received iC9.IL15.IL18 CAR-NKTs three days after administration of CID or vehicle. *n*=3, ** *p* < 0.01, unpaired two-tailed t-test. **D.** Serum cytokine quantification in indicated groups of mice, *n=*3 mice per group, mean (SD), * *p* < 0.05, ** *p* < 0.01, *** *p* < 0.001, **** *p* < 0.0001, ordinary one-way ANOVA of log_10_ transformed data.

**Supplementary Figure 4. IL-18 mediates superior antitumor activity in a NB model with intermediate GD2 expression.**

**A.** *In vivo* tumor challenge experimental design. NSG mice were injected via tail vein with luciferase-expressing CHLA136 NB cells, and CAR-NKTs were infused through the same route seven days later. Tumor growth was assessed weekly by bioluminescence imaging, and weight was measured every other day. **B.** CHLA136 tumor progression was measured over seven weeks by bioluminescence imaging after administration of the indicated CAR-NKTs. (*n*=4-10 mice per group) **C.** Quantification of bioluminescence signal for mice shown in (**B**). **D.** Percent weight change for mice shown in (**B**) after NKT infusion. **E.** Kaplan-Meier survival curve for mice in (**B**). *** *p* < 0.001, **** *p* < 0.0001, Log-rank (Mantel-Cox).

**Supplementary Figure 5. Evaluation of gene expression changes in iC9.IL15 and iC9.IL18 expressing CAR-NKTs**

**A.** Experimental design for RNAseq studies. RNA was isolated from CAR-NKTs prior to stimulation for baseline assessment. Cells were then stimulated one or three times with CHLA255 cells, dead cells were removed, and bulk RNAseq performed. scRNAseq was performed after one stimulation only. Volcano plot depicting differentially expressed genes in CAR-NKTs expressing IL18 versus IL15 before stimulation (**B**), after one tumor stimulation (**C**), and after three tumor stimulations (**D**). **E.** Representative flow cytometry plots for the CD25 and CD86 expression in CAR^+^ NKT cells (top) and summary data for CD86 and CD25 expression in CAR-NKTs (bottom) before tumor stimulation. Mean shown, *n*=8 donors in two independent experiments. * *p*<0.05, ** *p*<0.01, paired two-tailed t-test. **F.** Significantly enriched GSEA terms in CAR-NKTs expressing iC9.IL18 versus iC9.IL15 after three tumor stimulations.

**Supplementary Figure 6.**

**A.** Heatmap of top differentially expressed genes in clusters depicted in Fig. 5A.

**Supplementary Figure 7. IL-18 broadly enhances CAR-NKT metabolism**

**A-C.** MitoTracker Green (MG), tetramethylrhodamine ethyl ester perchlorate (TMRE), or BODIPY staining was performed at baseline and/or after one tumor stimulation to evaluate metabolic parameters in CAR-NKTs by flow cytometry. **A.** Flow cytometry histograms depicting MG (left) and TMRE (right) staining in CAR^+^ NKT cells from a representative donor at baseline. **B.** Summary data for MG and TMRE staining after tumor stimulation (left; mean, *n*=12 donors, three independent experiments) and TMRE-to-MG ratio calculated from values in left panel (right; mean, *n*=12 donors, three independent experiments), *** *p* < 0.001, **** *p* < 0.0001, paired two-tailed t-test. **C.** Representative histogram depicting BODIPY uptake in unstimulated CAR^+^ NKTs transduced with iC9.IL15 or iC9.IL18 (left). Summary data of fatty acid uptake measured using BODIPY in CAR^+^ NKTs before and after tumor stimulation. *n*=12 donors, mean, ** *p* < 0.0001, **** *p* < 0.001, paired two-tailed t-test (right). **D.** Significant differentially expressed metabolites in CAR-NKTs expressing iC9.IL18 versus iC9.IL15 at baseline (*n*=5 donors, *p<0.05*). **E.** Representative histogram depicting 2-NBDG uptake in unstimulated CAR^+^ NKTs transduced with iC9.IL15 or iC9.IL18 (left). Summary of 2-NBDG uptake measured by flow cytometry in CAR^+^ NKTs before and after tumor stimulation. *n*=12 donors in three independent experiments. Mean, ns=not significant, **** *p* < 0.0001, paired two-tailed t-test (right). **F.** GSEA analysis of oxidative phosphorylation (left) and glycolysis (right) gene signatures in CAR-NKTs expressing iC9.IL18 versus iC9.IL15 at baseline. **G.** Oxidative phosphorylation and glycolysis signatures in scRNAseq data set shown in Fig. 5A.

**Supplementary Figure 8. IL-18 increases glutaminolysis and purine metabolism in CAR-NKT cells**

**A.** Pathway enrichment analysis of DEMs in **Fig. S7E** before stimulation **B.** Pathway enrichment analysis of DEMs in Fig. 6C after stimulation. **C.** Normalized glutamine counts in CAR-NKTs before stimulation. n=5 donors, mean, *** *p*<0.001, Wald test (left). Normalized glutamine counts in CAR-NKTs after stimulation. n=4 donors, mean, * *p*<0.05, Wald test (right). **D.** Heatmap depicting glutaminolysis related genes before and after stimulation. **E.** Heatmap depicting purine metabolism related genes before and after stimulation.

