## Supplemental Figures for "IL-18 metabolically reprograms CAR-expressing natural killer T cells and enhances their antitumor activity against neuroblastoma"

**Supplementary Figure 1.** Generation and characterization of NKT cells co-transduced with a GD2.CAR and an iC9.cytokine construct

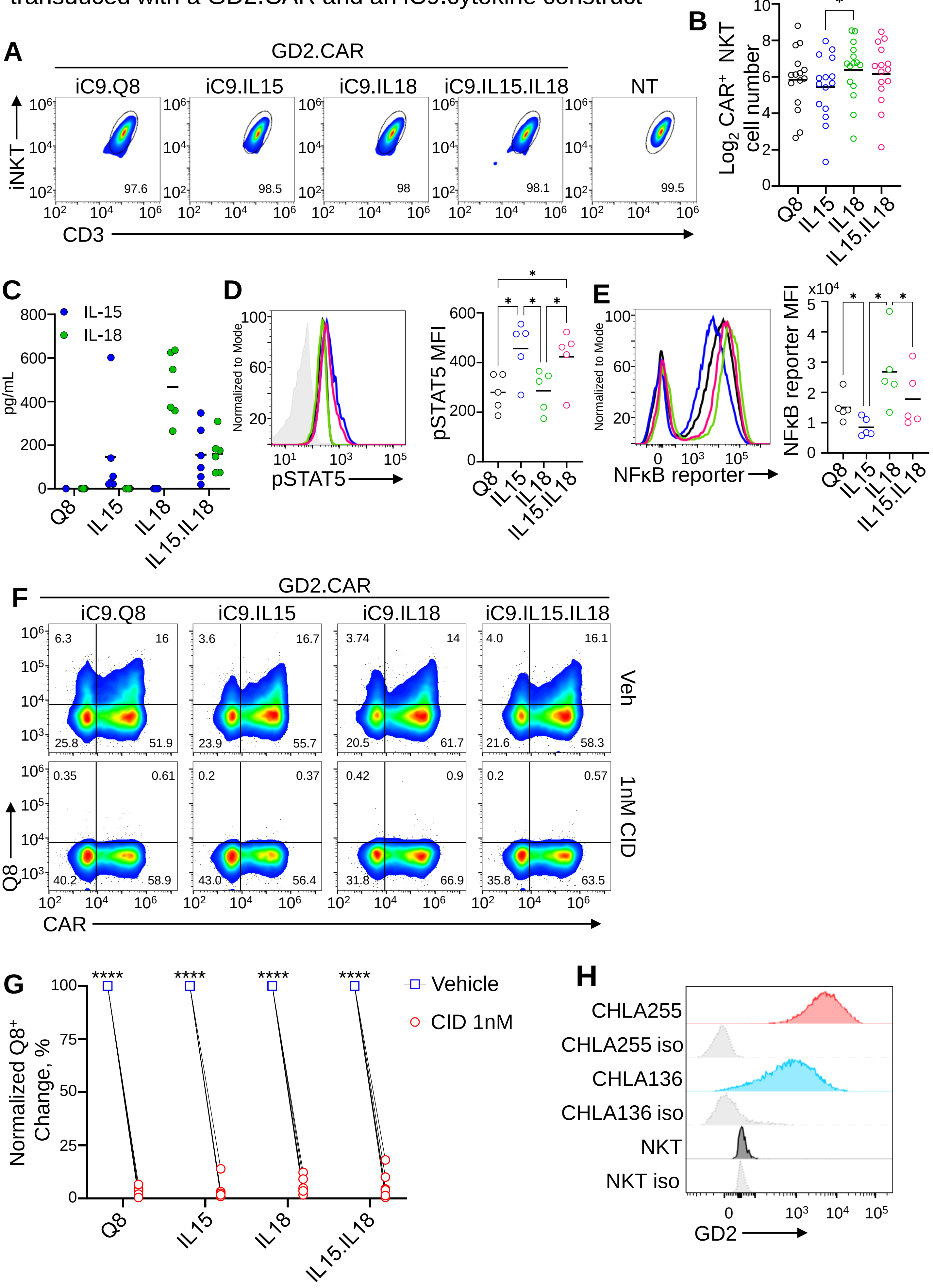

**Supplementary Figure 2.** iC9.IL18 expression enhances CAR-NKT function after repeated tumor challenge

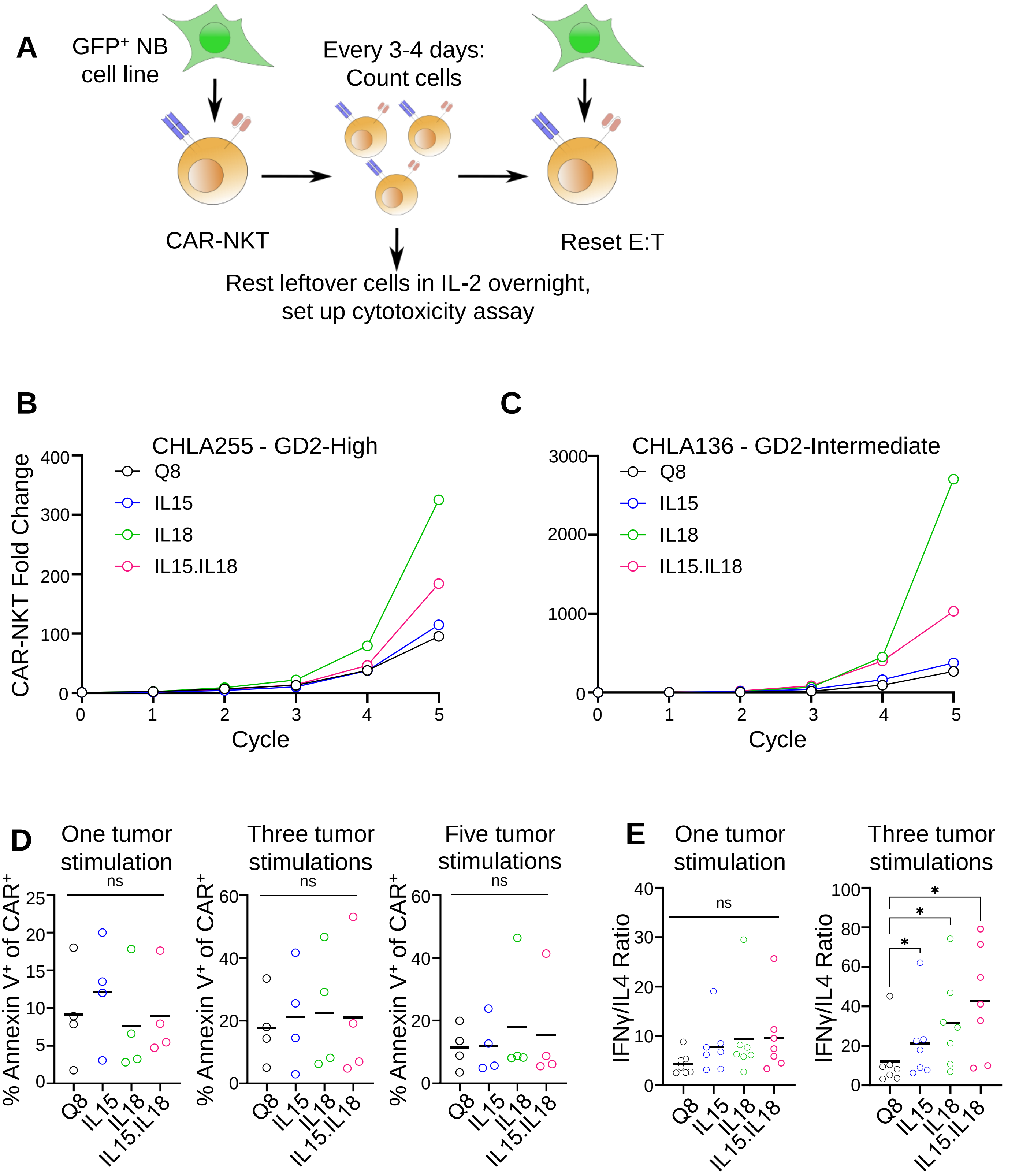

**Supplementary Figure 3.** Characterization of iC9.IL15.IL18 toxicities in mice

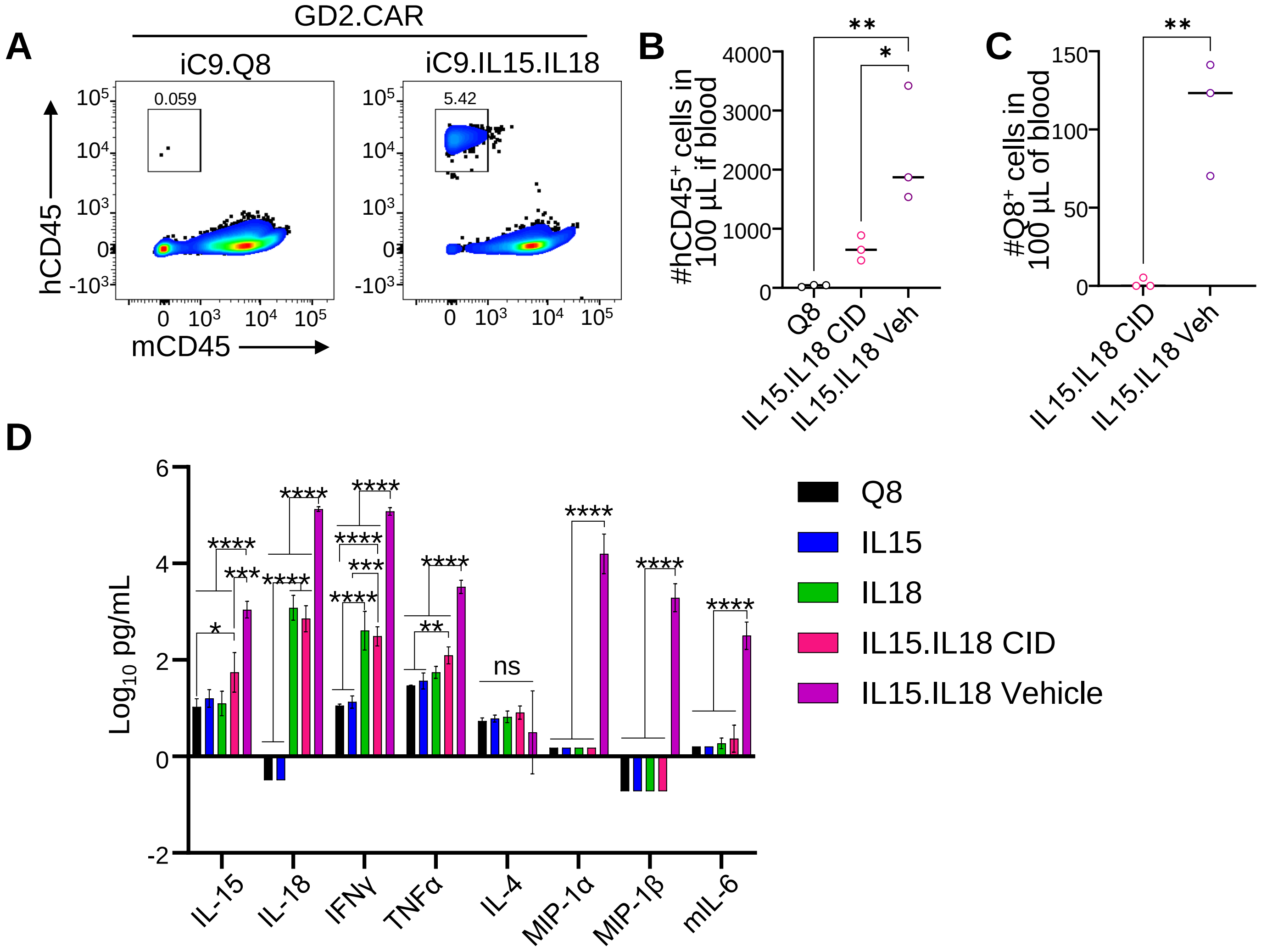

**Supplementary Figure 4.** IL-18 mediates superior antitumor activity in a NB model with intermediate GD2 expression

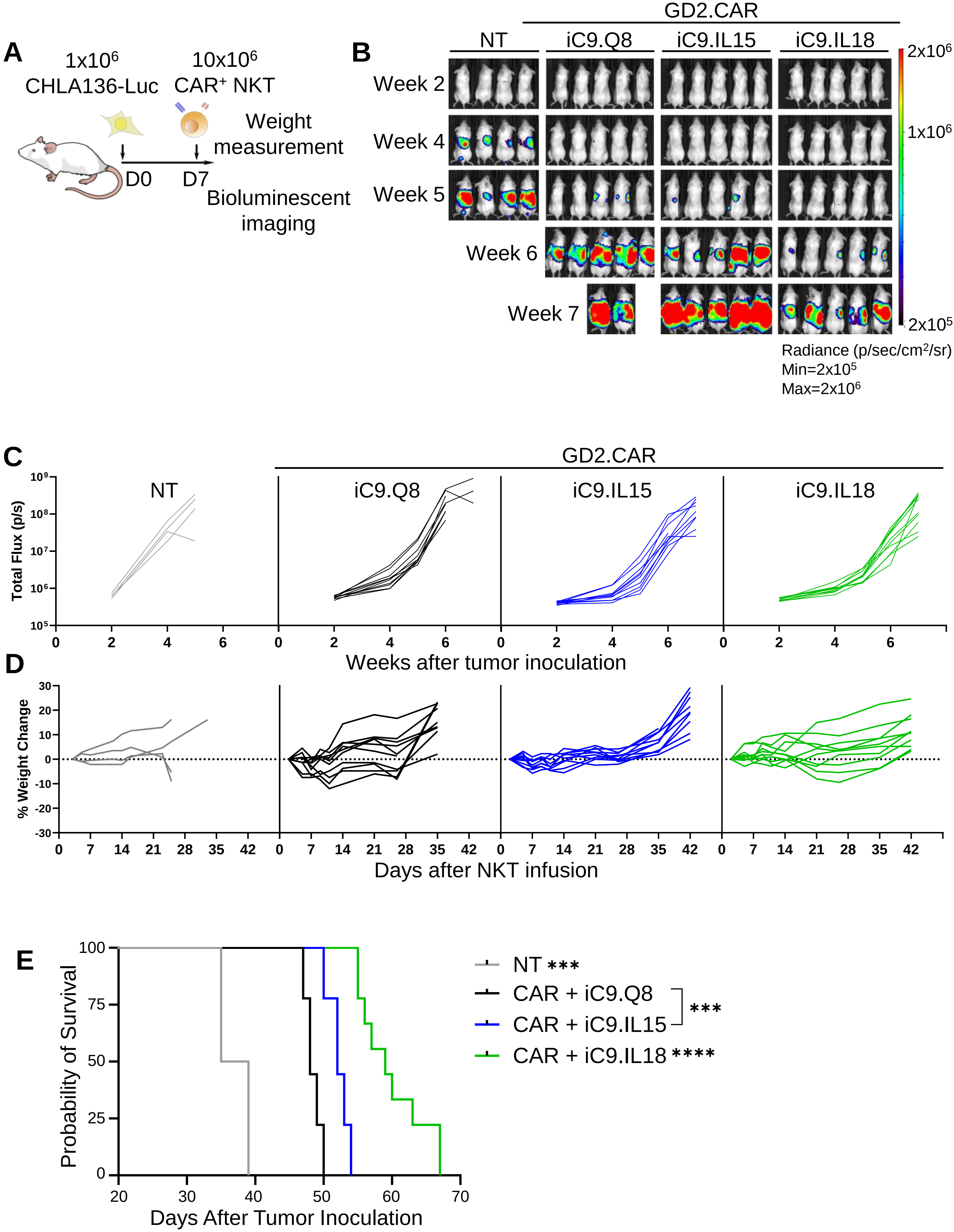

**Supplementary Figure 5.** Evaluation of gene expression changes in iC9.IL15 and iC9.IL18 expressing CAR-NKTs

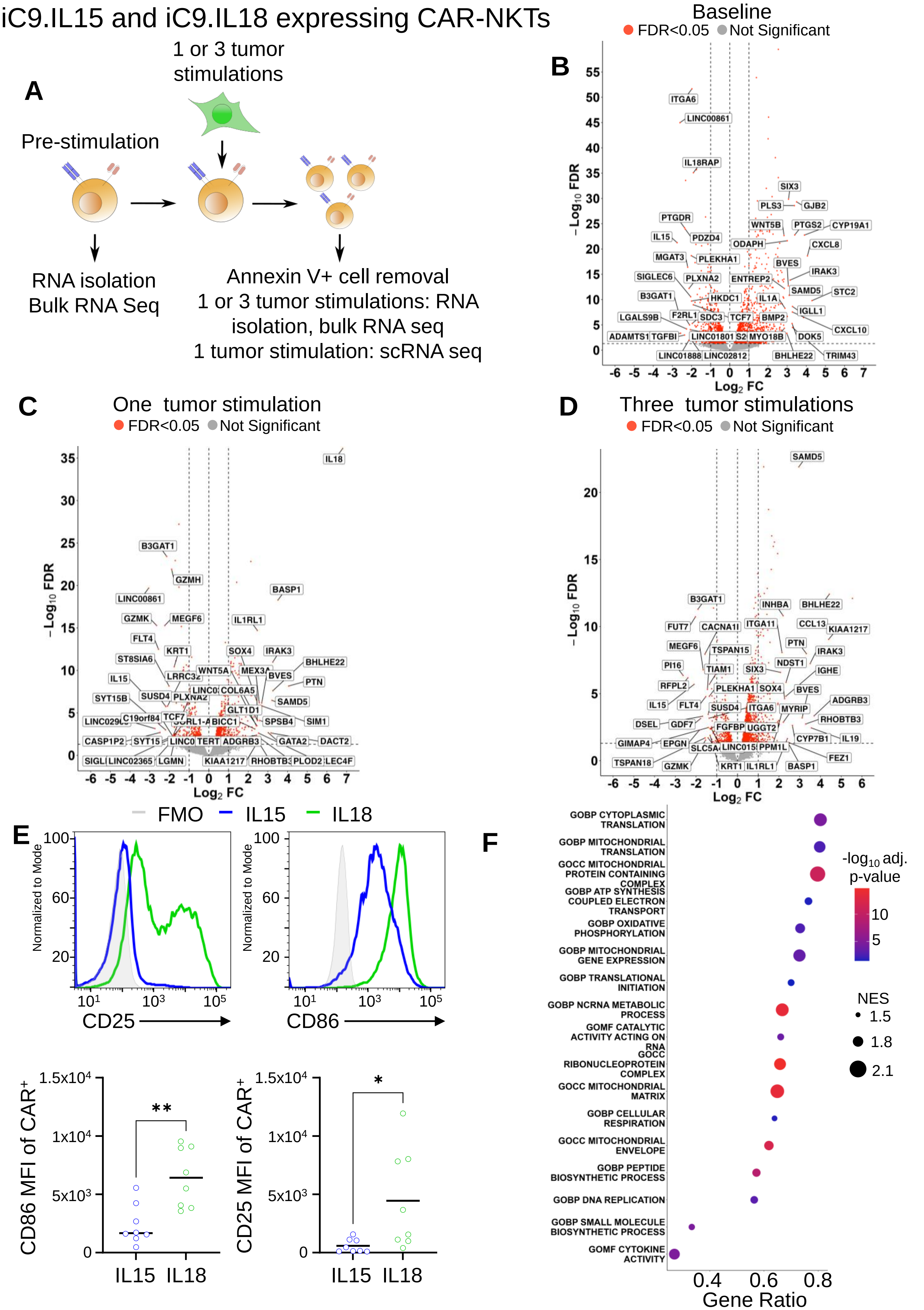

### Supplementary Figure 6.

**A.**

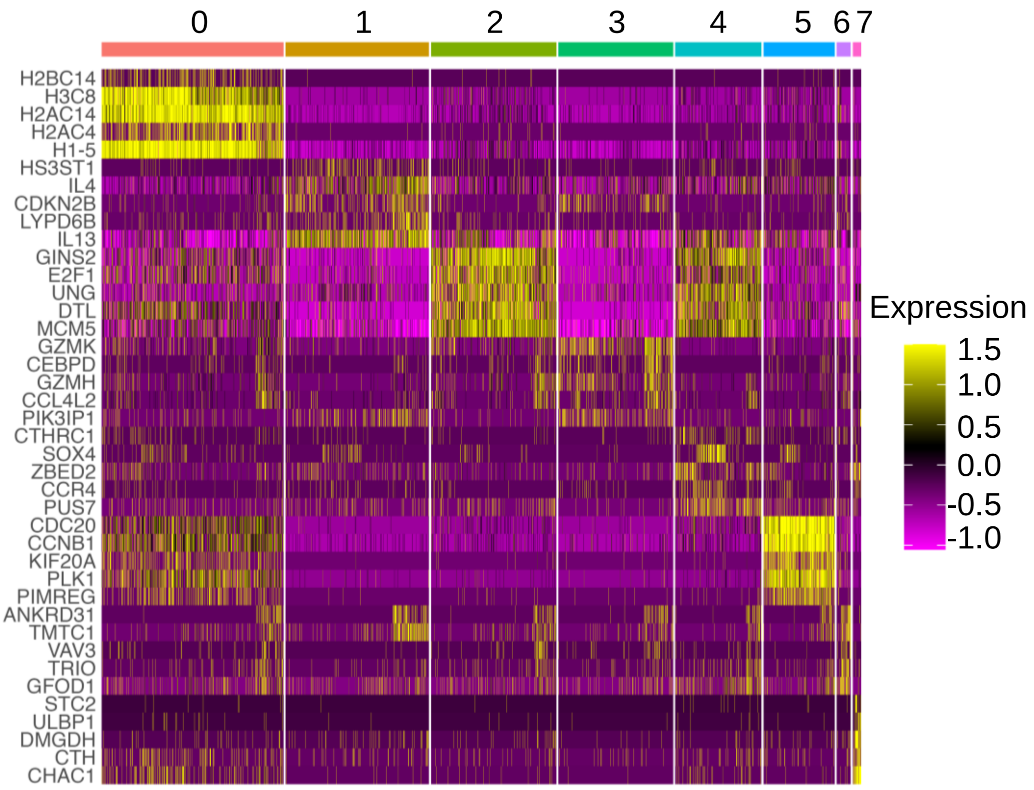

**Supplementary Fig. 7** IL-18 broadly enhances CAR-NKT metabolism

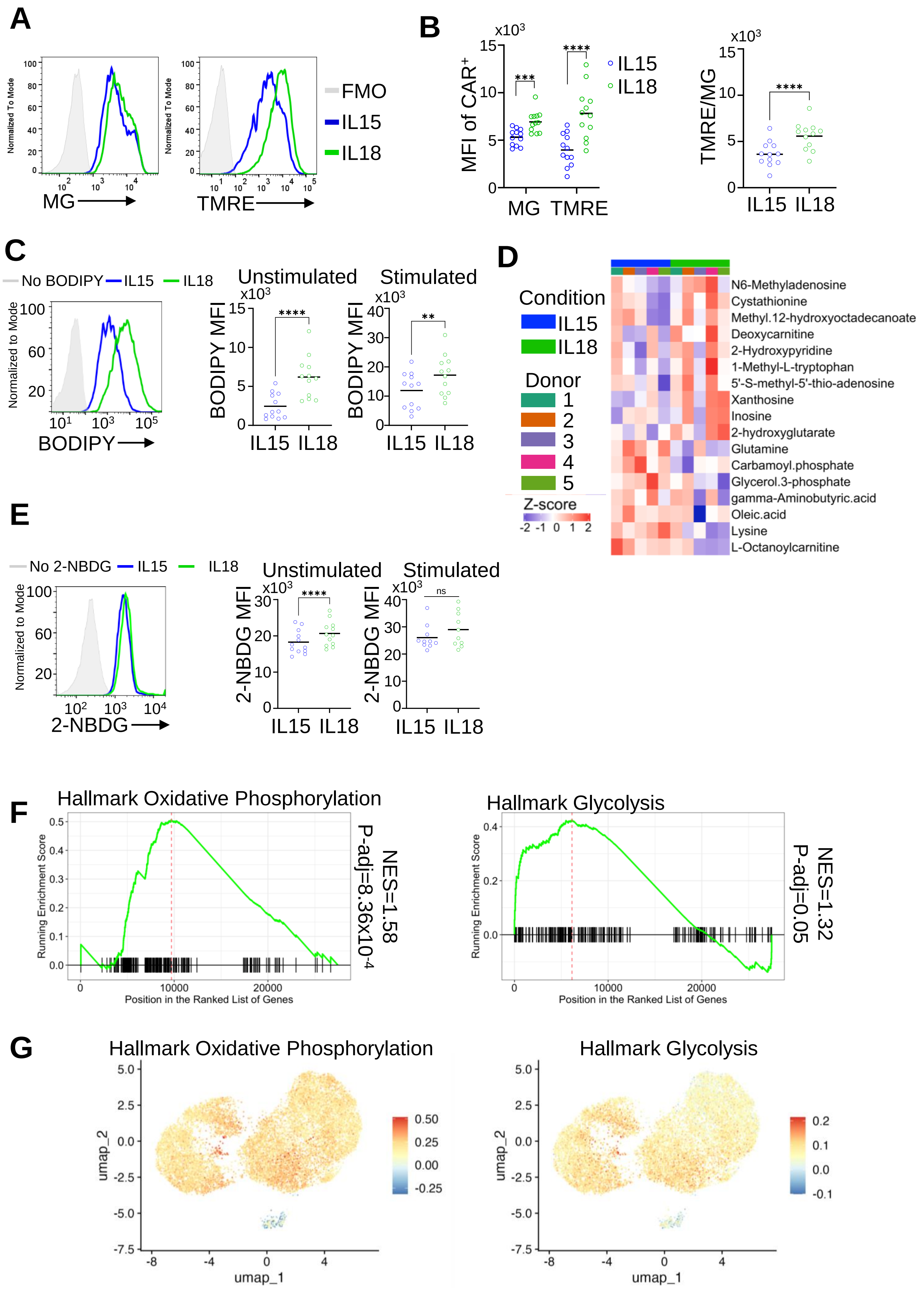

**Supplementary Figure 8**. IL-18 increases glutaminolysis and purine metabolism in CAR-NKT cells

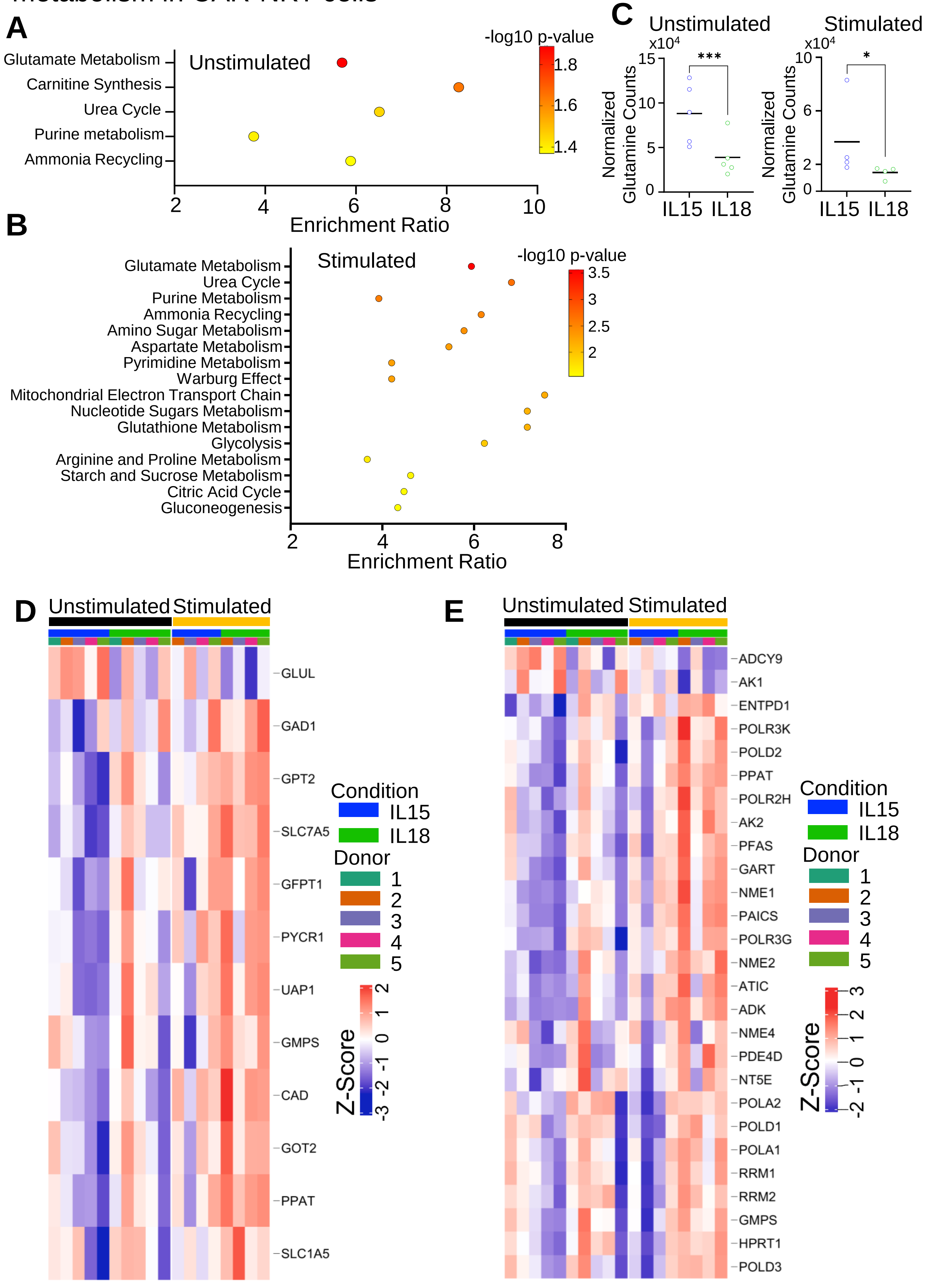
